# Development of a humanized anti-FABP4 monoclonal antibody for treatment of breast cancer

**DOI:** 10.1101/2024.05.12.593748

**Authors:** Jiaqing Hao, Rong Jin, Yanmei Yi, Xingshan Jiang, Jianyu Yu, Zhen Xu, Nicholas J. Schnicker, Michael S. Chimenti, Sonia L. Sugg, Bing Li

**Author notes:** **Corresponding Author** Bing Li, Professor at Department of Pathology, University of Iowa. Address: 431 Newton Road, Iowa City, IA, 52242, USA,. These authors contribute equally to the paper (Jiaqing Hao, Rong Jin, Yanmei Yi).

## Abstract

**Background:** Breast cancer, lung cancer, and colorectal cancer are the primary contributors to newly diagnosed cases among women, with breast cancer representing the second highest proportion of the total. The treatment protocols vary depends on different stages of breast cancer, and numerous clinical trials are ongoing based on the data derived from laboratory. Our studies demonstrate that circulating adipose fatty acid binding protein (A-FABP, or FABP4) links obesity-induced dysregulated lipid metabolism and breast cancer risk, thus offering a new target for breast cancer treatment.

**Methods:** We immunized FABP4 knockout mice with recombinant human FABP4 and screened hybridoma clones with specific binding to FABP4. The potential effects of antibodies on breast cancer cells *in vitro* were evaluated using migration, invasion, and limit dilution assays. Tumor progression *in vivo* was evaluated in various types of tumorigenesis models including C57BL/6 mice, Balb/c mice, and SCID mice. The phenotype and function of immune cells in tumor microenvironment were characterized with multi-color flow cytometry. Tumor stemness was detected by ALDH assays. To characterize antigen-antibody binding capacity, we determined the dissociation constant of S-V9 against FABP4 via surface plasmon resonance. Further analyses in tumor tissue were performed using 10X Genomics Visium spatial single cell technology.

**Results:** Herein, we report the generation of humanized monoclonal antibodies blocking FABP4 activity for breast cancer treatment in mouse models. One clone, named 12G2, which significantly reduced circulating levels of FABP4 and inhibited mammary tumor growth, was selected for further characterization. After confirming the therapeutic efficacy of the chimeric 12G2 monoclonal antibody consisting of mouse variable regions and human IgG1 constant regions, 16 humanized 12G2 monoclonal antibody variants were generated by grafting its complementary determining regions to selected human germline sequences. Humanized V9 monoclonal antibody showed consistent results in inhibiting mammary tumor growth and metastasis by affecting tumor cell mitochondrial metabolism.

**Conclusions:** Our current evidence suggest that targeting FABP4 with humanized monoclonal antibodies represents a novel strategy for the treatment of breast cancer and possibly other obesity- associated diseases.

## Background

Surpassing lung cancer for the first time in 2020, breast cancer has become the most common cancer in women diagnosed in the U.S. and worldwide(1). The incidence of breast cancer has increased dramatically from 641,000 cases in 1980 to more than 2.3 million in 2020(1, 2). Despite new treatment options in improving survival, around 685,000 women still die of breast cancer worldwide annually(1). Multiple risk factors, including genetic mutations and background, family and reproductive history, aging and exposures to toxins, contribute to breast cancer risk. However, the global rates of breast cancer incidence and mortality have continuously increased in the past few decades(3), suggesting that other etiological factors besides the above-mentioned factors contribute to the increasing rates of breast cancer.

Obesity is a complex condition with multiple contributing factors, including genetics, environment, behavior, and metabolism(4, 5). In modern society, owing to excessive calorie intake and a sedentary lifestyle(6, 7), obesity has risen at an alarming rate in the U.S. According to CDC statistics, adult obesity prevalence reached 42.4% in 2017-2018. Nearly half of Americans are projected to become obese by 2030. As a result, extra energy stored as lipids in various cells and tissues can negatively affect cell metabolism and homeostasis. In obese patients, adipose tissue contains crown-like structures formed by hypertrophic adipocytes surrounded by macrophages(8). The interaction between macrophages and adipocytes promotes obesity-associated chronic inflammation and further pathological alterations. As an endocrine organ, adipose tissue produces multiple mediators (*e.g.,* adipokines, cytokines, chemokines and hormones) to maintain metabolic balance. Dysregulation of these mediators leads to obesity-associated diseases, including at least 13 types of cancer(9). Epidemiologic studies have confirmed that obesity increases not only the risk of breast cancer in postmenopausal women, but also the mortality from breast cancer in women of all ages(10–12).

Given the prevalence of obesity and increased risk of breast cancer in obese patients, several cellular and molecular mechanisms have been proposed to explain the obesity/cancer axis, which includes cancer-associated adipocytes, obesity-related inflammatory cytokines (*e.g.,*IL-6 and TNFα), lipids (*e.g.*, lysophosphatidic acid and prostaglandins), adipokines (*e.g.,* leptin and adiponectin), insulin/insulin-like growth factors (IGFs), and sex hormones(9, 13, 14). Although substantiated by significant clinical and experimental data, these mechanisms remain contentious due to the complex multisystem interactions between obesity and cancer(15). In exploring obesity/breast cancer risk, we have identified adipose fatty-acid binding protein (A-FABP, also known as FABP4, aP2) as a new molecular mechanism linking obesity-associated breast cancer development(16–18). Traditionally recognized as an intracellular lipid chaperone, FABP4 is mainly expressed in adipose tissue, facilitating fatty-acid transportation, metabolism and responses(19, 20). However, we demonstrate that obesity elevates FABP4 secretion from adipose tissue into the circulation, where extracellular FABP4 can directly target breast cancer cells, enhancing IL-6/STAT3/ALDH1-mediated tumor stemness and aggressiveness(16, 17). Circulating FABP4 bridges tumor-associated stromal cells to tumor stem cells and integrates adipokines to tumor-promoting signaling and lipid metabolism, thereby representing a new molecular link underlying obesity-associated breast cancer risk and mortality. Therefore, targeting circulating FABP4 represents a novel strategy for treatment of breast cancer.

In the current study, we immunized mice with recombinant FABP4 protein and screened over 1200 clones. We identified one clone, 12G2, which was able to effectively block FABP4 activity and inhibit mammary tumor growth in different mouse models. After evaluating the efficacy of its chimeric version and humanized variants, we successfully generated a humanized FABP4 monoclonal antibody (mAb), which offers great potential for treating breast cancer in the clinic.

## Materials and methods

### Generation of mouse monoclonal antibody

FABP4 mAbs were generated by immunization of 7-week-old female FABP4 knockout mice with full length human recombinant FABP4 protein. Briefly, 50μg protein emulsified with complete Freund’s adjuvant (Sigma) was subcutaneously (s.c.) injected on the back of the mice. Mice were boosted with 25μg protein mixed with incomplete Freund’s adjuvant (IFA) by s.c. at day 14 and day 28. The final boosting was conducted at day 50 with 25μg protein mixed with IFA by intravenous injection. Blood from immunized mice was collected for measurement of anti-FABP4 antibodies by ELISA. Mice with the high titer of anti-FABP4 antibodies were selected for splenocyte collection and fusion with Sp2/0 myeloma cells (ATCC). Hybridoma generation was performed using the ClonalCell^TM^-HY kit (STEMCELL Technologies). Of note, compared to conventional hybridoma selection and cloning, this method uses a methylcellulose-based semi-solid medium, which increases the diversity of clones that can be easily identified and isolated, enabling hybridoma selection and cloning to complete in a single step(21). On day 12 after fusion, a total of 1248 single clones in the semi-solid medium were collected and cultured for another 4-7 days. The supernatants were screened for reactivity to human recombinant FABP4 or FABP5 by ELISA. A total of 141 positive clones to FABP4 were selected for further screening with other sources of human or mouse FABP4 protein (Cayman Chemical). Finally, 25 positive clones with specific reactivity to both human and mouse FABP4, but not FABP5, were identified. These which were able to produce ample ascites and with better affinity to FABP4 were selected for antibody production.

### Antibody purification from ascites

Ascites of the selected clones of hybridoma cells were developed in 8-week-old female Balb/c mice (n=6-7 mice/group). Briefly, 0.5ml pristane was intraperitoneally (i.p.) injected into each mouse. After pristane priming for 7-10 days, 5×10^6^ hybridoma cells in 400 *μ*l PBS were i.p. injected into each mouse. Ascites developed 5-7 days after hybridoma cell injection were harvested from the second week using the 19-gauge needles. Cellular components in the ascites were removed by centrifugation at 2000 rpm for 15 mins. For monoclonal antibody purification, ascites was diluted by adding 4 volumes of 60mM acetate buffer with a final pH of 4.5. Caprylic acid was dropwise added to the diluted ascites at 25 μl/ml. The mixture was stirred for 30 mins and then centrifuged for 30 min at 10000g. The supernatant was collected and mixed with 1/10 volume 10x PBS after nylon mesh filtration. After pH adjustment to 7.4, ammonium sulfate (0.277g/ml) was slowly added to the solution at 4 °C and stirred for additional 30 mins before centrifugation for 30 min at 5000g. The precipitated antibody was resuspended in small volume of PBS. The purity and quantification of different monoclonal antibodies were determined for further applications.

### Evaluation of antibody therapeutic efficacy using breast cancer mouse modes

Mouse models of breast cancer were used to evaluate the potential therapeutic efficacy of different clones of purified FABP4 antibodies. Mouse experiments were performed according to the approved procedures by the Institutional Animal Care and Use Committee (IACUC) at the University of Iowa. C57BL/6-derived mammary tumor cells E0771 (5×10^5^cells/mouse) were orthotopically injected into mammary fats of C57BL/6 mice (8-10 weeks old). After tumor injection, mice were randomly divided into several groups and treated with different clones of purified antibodies (5mg-30mg/kg, twice/week). Mice treated with same volume of PBS were used as controls. When the tumors were palpable, the length and width of the tumors were measured by a caliper every three days. The volume of the tumors was calculated using the formular of 0.5 x length x width^2^ as descripted before(22, 23). To confirm the therapeutic efficacy of the selected clone of anti-FABP4 antibody, other breast cancer models, including Balb/c-derived MMT, 4T1 models, human-derived MCF-7 breast cancer in SCID mice, were also used in the present studies.

### Production of chimeric and humanized anti-FABP4 antibodies

For chimeric antibodies, anti-FABP4 hybridoma clones (e.g. 12G2, 6H10) were sequenced and DNA sequences of the VH and VL regions were identified. Recombinant chimeric antibodies consisting of mouse VH and VL and human IgG1 constant regions were expressed and purified in HEK293 cells. For humanized antibody production, parental VH and VL sequences were run through a CDR grafting algorithm to transfer the CDRs from the original framework onto the most matched human germline sequences. To ensure that no highly undesirable sequence liabilities were introduced into the humanized sequences, identified high-risk motifs were removed through mutagenesis. A total of 16 antibody variants combined of 4 humanized heavy chains and 4 light chains were generated using HEK293 mammalian cells. Purified antibodies were analyzed for purity by SDS-PAGE and their concentrations were determined by UV spectroscopy.

### Aldehyde dehydrogenase (ALDH) assay

The ALDEFLUOR kit (Cat. #01700 from STEMCELL technologies) was used to detect ALDH activity for both tumor tissues and tumor cell lines. To obtain single cells from tumor tissues, E0771 and MCF7 tumors were removed from euthanized mice and mechanically dissociated into smaller fragments. These fragments were then treated with 6 mL tri-enzyme solution containing collagenase type 2, hyaluronidase, and DNase I in RPMI-1640 medium containing 5% FBS and incubated at 37°C for 45 minutes on an orbital shaker at speed of 50 rpm. Following enzymatic digestion, the cell suspensions were collected by vertexing, filtration, removal of tri-enzyme solution, and two washes with cold 1x PBS. The detection of ALDH activity from both single-cell suspensions derived from tumor tissues and tumor cell lines were followed the protocol provided by the manufacturer’s instruction of the ALDEFLUOR kit.

### Tumor migration, tumor invasion, and limit dilution assays

To assess the blocking activity of anti-FABP4 antibodies, individual antibodies (1 µg/mL) and human FABP4 protein (200ng/mL) were mixed for 15 minutes before performing following assay. FABP4 and PBS alone were served as controls. 1) wound-healing migration was performed to assess whether antibody was able to inhibit FABP4-mediated tumor cell migration. To induce a linear wound in the cellular monolayer, the confluent cells were mechanically scratched using a 200 µL plastic pipette tip in a six well-plate containing 2.5 mL cell culture medium. Subsequently, the scratched monolayer was carefully washed with pre-warmed 1 x PBS to eliminate any debris. Following a 96-hour incubation period at 37°C, the migration of cells towards the wound site was captured using light microscopy, and the migration distance was quantified using Image J software. 2) For tumor invasion assay, MCF-7 cells were cultured to form spheres in hanging drops of culture medium on the lid of cell culture dishes as previously described(17). Briefly, following a seven-day incubation period, the spheroids from the lid were transferred into an equivalent volume and combined with rat tail type I collagen, reaching a final concentration of 1.7 mg/mL. This mixture was then embedded in a 24-well plate to establish a 3D culture system. FABP4/antibody mixture, FABP4 protein and control PBS were added into 1 mL cell culture medium, respectively. Quantitative analyses were performed by measuring the maximal invasive distance (longest distance from the spheroid radius) and the invaded area (total invaded area minus the spheroid area) using Image J software. 3) For in vitro limiting dilution assay (LDA), tumor cells were serially diluted to obtain cell concentration at a range of 1000, 500, 250, 125, 62, 31, 16, 8 cells and seeded into an ultra-low attachment 96-well plate containing 200 µL cell culture medium, exposed to FABP4-antibody mixture, FABP4 protein and control PBS for a duration of 48 - 96 hours. Subsequently, cell spheres were determined using microscope and calculated the cancer cells initiating frequency and significance using online software (Extreme limiting dilution analysis, ELDA @ http://bioinf.wehi.edu.au/software/elda/index.html) following the methodology outlined by Hu and Smyth(24).

### Characterization of antibody/antigen binding

The binding of anti-FABP4 antibodies with FABP4 was evaluated by ELISA and Biocore assays. For ELISA, FABP4 protein was diluted with 1 x PBS and coated to a 96-well plate at a final volume of 100 µL overnight. After blocking with 5% BSA at room temperature for 1hour, anti-FABP4 antibodies (e.g. chimeric, humanized) were diluted with 5% BSA solution and added into indicated wells. The plate was washed three times using 200 µL of 1 x PBS containing 0.5 % Tween-20 and incubated with 100 µL of secondary antibody solution containing goat anti-human IgG conjugated with HRP (Cat. #A18805 from Invitrogen) at a dilution of 1:10000 in 5% BSA solution for 1hour. Color development was performed by adding 100 µL of substrate solution and incubating for 5 minutes at room temperature before the reaction was stopped by 100 µL of 2N sulfuric acid. OD value was acquired using a BioTek Synergy LX Multimode Reader. For binding affinity measurement, FABP4 (10μg/ml) was immobilized on CM5 sensor chip of Biocore 8K using maleimide EDC/NHS coupling. Antibody (V9) at a defined concentration is flown over CM5 chip and response captured over time, showing the progress of the interaction and association/dissociation cycle. After different concentrations are successively tested, the kinetics parameters and affinity are calculated using BIA-evaluation software.

### Immunophenotype analysis by flow cytometry

Immune phenotypes were performed using multi-color staining panel designed by improved version of full spectrum viewer in Cytek Cloud. Signle-cell suspension of the primary tumor was resuspended in 1 x PBS containing 0.5% FBS and kept in the ice all the time. Cells were pre-incubated with anti-mouse CD16/CD32 antibody (Cat. # 101302 from Biolegend) for 5 minutes to block Fc receptors. Surface staining were prepared using the following antibodies: Zombie-violet (Cat. #423108, Biolegend), anti-mouse CD45 (Cat. #103116, Biolegend), anti-mouse CD11b (Cat. #612800, BD), anti-mouse CD3 (Cat. #100355, Biolegend), anti-mouse CD4 (Cat. #740208,BD), anti-mouse CD8 (Cat. #752642, BD), anti-mouse F4/80 (Cat. #123120, Biolegend), anti-mouse MHCII (Cat. #107604, Biolegend), anti-mouse Ly6G (Cat. #127664, Biolegend), anti-mouse NK1.1 (Cat. #108736, Biolegend), anti-mouse B220 (Cat. #103232, Biolegend), and anti-mouse CD11c (Cat. #117349, Biolegend). The intracellular cytokines staining with anti-mouse IL-6 (Cat. #504508, Biolegend) and anti-mouse TNFα (Cat. #506338, Biolegend) were fixed and permeabilized using True-Nuclear transcription factor buffer set (Cat. #424401 from Biolegend) according to the manufacturer’s introduction. All samples were acquired with an Cytek Aurora instrument. Data were analyzed with FlowJo (BD).

### Spatial transcriptomics and analysis

Fresh tumor tissues were removed and placed in a petri-dish and embedded with room temperature OCT without any bubbles on the tissue’s surface. The embedded samples were transferred into the cryo mold. The cryo mold containing OCT-embedded samples were put into the metal beaker with 2-methylbutane in a dewar of liquid nitrogen until the OCT was solidified. Sample cryosectioning, affixment to cDNA capture slide, H&E staining, tissue permeabilization, RNA capture, and cDNA synthesis were performed according to the 10 x Genomics Visium spatial transcriptomics’ methods.

The four visium libraries (two PBS tumors, and two S-V9-treated tumors) were sequenced on an Illumina NovaSeq 6000 located in the Iowa Institute of Human Genetics (IIHG) Genomics division. Paired-end reads were demultiplexed and converted from the native Illumina BCL format to fastq format using an in-house python wrapper to Illumina’s ‘bcl2fastq’ conversion utility. The data were deposited to GEO repository (GSE264099). Bioinformatic analysis was carried out by the IIHG Bioinformatics division. Fastq data were merged across lanes and the PE reads were used as input for the 10X SpaceRanger pipeline in ‘count’ mode (v1.3.1). SpaceRanger was run on the Argon High-Performance Computing (HPC) cluster at the University of Iowa using 32 cores and 128GB of RAM per sample. The reference transcriptome was specified as ‘mm10-2020-A’ and the chemistry was specified as ‘Spatial 3’ v1’. QC analysis of the four samples showed no quality problems for each sample other than an alert that “Low Fraction Reads in Spots” was detected for 3 of 4 samples. Filtered barcode matrices were used as input for downstream analysis in Seurat (v5). Four Seurat objects were created from the individual barcode matrices and quality control (QC) metrics were visualized as violin plots that included the number of genes (nFeature), number of UMI (nCount) and percentage of mitochondrial UMI (percent_mt). We filtered cells that have less 100 features (low-quality cells). The filtered datasets were subjected to normalization, detection of variable features, scaling/centering and PCA analysis. Following this, the sample layers were integrated together using the “RPCA” method of integration available in Seurat 5 (https://satijalab.org/seurat/articles/integration_rpca). To cluster the cells, we used K-nearest neighbors (KNN) networks based on the calculated PCs. Modularity optimization was applied (Louvain method, resolution = 0.1) and a UMAP embedding was calculated. Searching for DEGs (cluster biomarkers), we found markers for every cluster compared with all remaining cells using the Wilcoxon Rank Sum test and a log_2_FC threshold of 0.4 and expressed in more than 30% of the cells. Cluster-wise DE analysis of the treatment effect of S-V9 vs PBS was carried out by using the “FindMarkers” function on the integrated object. Pathway analysis was carried out using g:Profiler (https://biit.cs.ut.ee/gprofiler/gost) and iPathwayGuide software (AdvaitaBio).

### H&E staining

Fresh tissues were obtained after removing primary tumor from euthanized mice. Lung samples were collected from the right inferior lobe and fixed immediately in 10% neutral buffered formalin. Air microbubbles were removed by placing lungs in a vacuum chamber for 5 minutes, then re-fixed the lungs into the fresh 10% neutral buffered formalin for 24 hours. Following serial alcohol dehydration (50%, 75%, 95%, and 100%), the samples were embedded in paraffin. The paraffin-embedded samples were sliced into 8 µm sections and stained in the DRS-601 Auto Stainer with hematoxylin and eosin (H&E) for 1 minutes. Slides were mounted with VectaMount® Express Mounting Medium (vector laboratories, H-5700-60), and were scanned by slide scanner (Leica Aperio GT 450) for quantification analysis. The metastatic tumor number and area was analyzed by the SlideViewer 2.7.0.191696 software.

### Quantification of serum biomarkers

Serum samples were collected at the end point of tumor challenge mouse model. Samples from each mouse were divided into aliquots and preserved in a −80C freezer for future purposes. The quantification of Fabp4(Cat. #CY-8077, MBL), IL-6(Cat. #431301, Biolegend) and glucose(Cat. #81692, Crystal Chem) levels were performed separately using ELISA kits in accordance with the instructions provided by the manufacturers.

### Statistical analysis

All data were presented as the mean ± SD unless notified specifically. For in vitro and in vivo experiments, a two-tailed, unpaired student t-test, were performed by GraphPad Prism 9. A statistics test is claimed to be significant if the p-value is less than 0.05.

## RESULTS

### Generation of anti-FABP4 monoclonal antibodies in mice

Human and mouse FABP4 share 91% amino acid sequence homology. To broaden the anti- FABP4 antibody epitope repertoire, we utilized FABP4 knockout mice and immunized them with recombinant human FABP4. Mice with high anti-FABP4 antibody titers were selected for hybridoma generation and clone selection (Figure S1A-S1C). After screening around 1248 clones in vitro, we identified at least 25 clones that specifically bound to FABP4 but not to FABP5 (Table S). Of these 25 clones, 6 clones were capable of inducing high-yield production of ascites (Table S).

To evaluate the potential neutralizing effect of these clones, we measured serum FABP4 levels in mice before and after ascites production. One clone, named 12G2, significantly reduced serum levels of FABP4 (Figure 1A). Interestingly, when MMT mammary tumor cells were implanted in mice with or without 12G2 ascites (Figure 1B), MMT tumor growth and weight in mice with 12G2 ascites were significantly reduced compared to mice without 12G2 ascites (Figure 1C, 1D). To verify the tumor inhibition effect, we purified antibodies from the 12G2 and two other clones (12H2 and 6H10) and used them to treat E0771, a commonly used breast cancer mouse model (Figure 1E). Compared to the 12H2 (Figure S1D) and 6H10 (Figure S1E) clones, the 12G2 clone significantly inhibited E0771 tumor growth (Figure 1F) and weight (Figure 1G). Moreover, serum levels of FABP4 (Figure 1H), IL-6 (Figure 1I), but not glucose (Figure 1J), were significantly reduced in response to 12G2 treatment. Altogether, these data suggest the 12G2 clone as a potential therapeutic monoclonal antibody targeting FABP4.

**Figure. 1.**
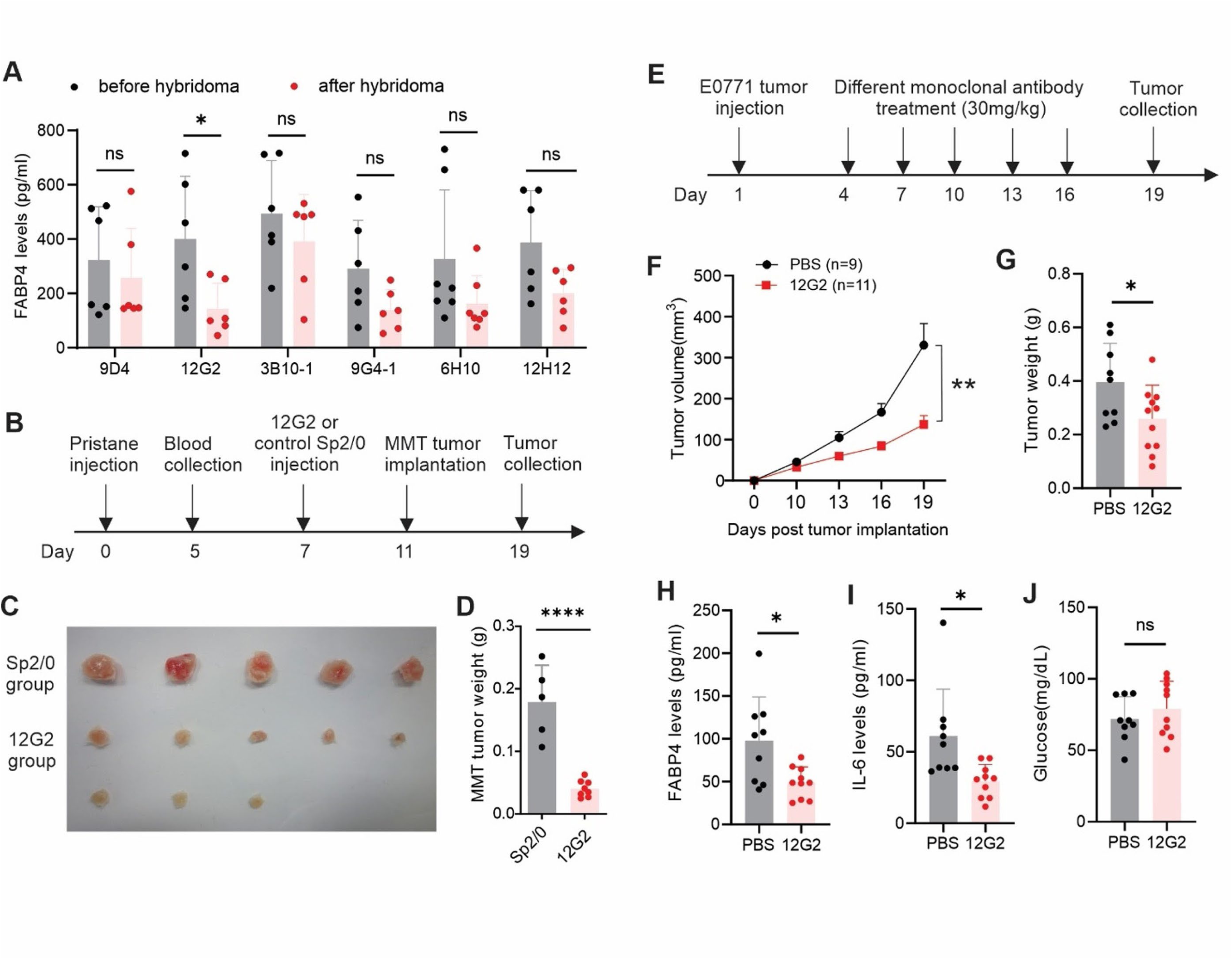
Screening of anti-FABP4 mAbs for treatment of mammary tumors. (A) Measurement of circulating FABP4 levels in mice before and after injection of different anti-FABP4 hybridoma clones by ELISA (*p<0.05, ns, non significant). (B) Schematic of evaluating the effect of anti- FABP4 hybridomas on MMT mammary tumor growth in vivo. (C, D) Tumor size (C) and weight (D) in mice injected with SP2/0 or 12G2 hybridoma, respectively (****p<0.0001). (E) Schematic of evaluating the effect of purified 12G2 antibody on E0771 mammary tumor growth in vivo. (F-G) E0771 tumor growth curve (F) and weight (G) in mice treated with 12G2 antibody or PBS control, respectively (*p<0.05, **p<0.01). (H-J) Serum levels of FABP4 (H), IL-6 (I) and glucose (J) in E0771 tumor-bearing mice treated with PBS or 12G2 antibody for 19 days (*p<0.05, ns, non-significant).

### Evaluation of the efficacy of chimeric mouse/human anti-FABP4 antibodies

To verify the therapeutic potential of the 12G2 clone, we generated chimeric mouse-human antibodies by joining the variable regions of 12G2 or 6H10 (as a control) to human IgG1 constant regions (Figure 2A). After the purification of the two chimeric antibodies (Figure S2A), we used an MCF-7 xenograft model to test their therapeutic efficacy in vivo (Figure 2B). Consistent with their parental clones, the chimeric 12G2 mAb exhibited better efficacy than 6H10 mAb in inhibiting MCF-7 tumor growth and size in SCID mice (Figure 3C, 3D). Treatment with the chimeric 12G2 antibody also significantly reduced MCF-7 tumor weight in SCID mice (Figure 3E). Interestingly, FABP4-mediated MCF-7 cell invasion and ALDH1 activity were significantly inhibited by the treatment of the chimeric 12G2 antibody (Figure 2F-2G, Figure S2B-S2D), further corroborating a specific role of 12G2 in blocking FABP4 activity in vitro. Moreover, using colony formation assays, we demonstrated that 12G2 significantly inhibited FABP4-induced colony-initiating cell frequency in different breast cancer tumor cell lines, including E0771, M158 and MCF7 (Figure S2E-S2G). Altogether, these *in vitro* and *in vivo* studies strongly supported the therapeutic efficacy of the chimeric 12G2 antibody.

**Figure 2.**
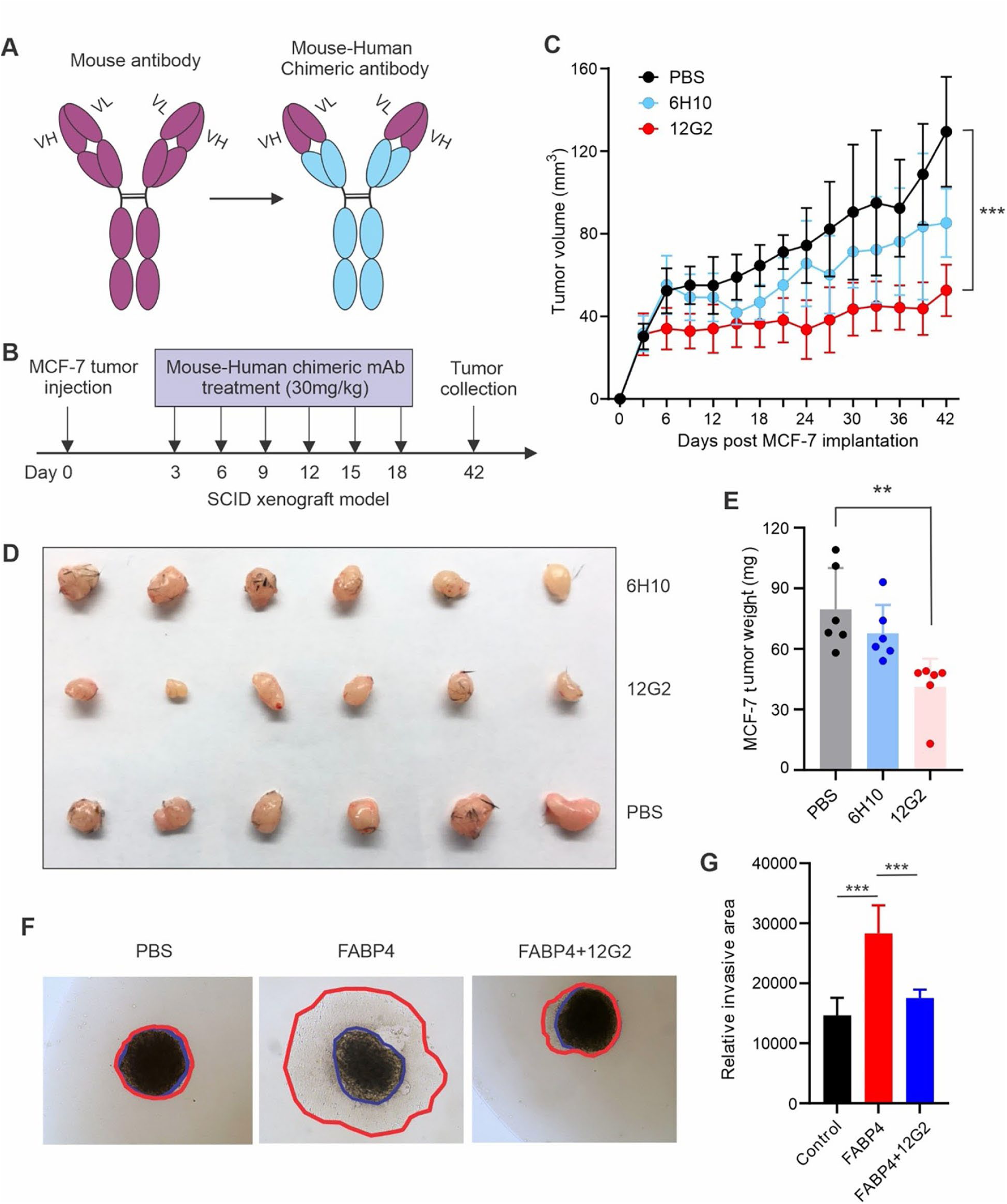
Evaluating therapeutic and blocking activity of chimeric anti-FABP4 antibodies. (A) Schematic of production of chimeric anti-FABP4 antibodies. (B) Schematic of evaluating the efficacy of chimeric antibody using MCF-7 breast cancer model in vivo. (C) Tumor growth curve of MCF-7 tumor in SCID mice treated with chimeric 6H10, 12G2 or PBS, respectively (***p<0.001). (D) Tumor size of MCF-7 in mice treated with 6H10, 12G2 or PBS, respectively, for 6 weeks. (E) Tumor weight of MCF-7 in mice treated with 6H10, 12G2 or PBS, respectively, for 6 weeks (**p<0.01). (F-G) Measurement of MCF-7 tumor cell invasion in the presence of PBS, FABP4 (100ng/ml), or FABP4+12G2 in vitro (F). Tumor invasion areas in each group are shown in panel G (***p<0.001).

**Figure 3.**
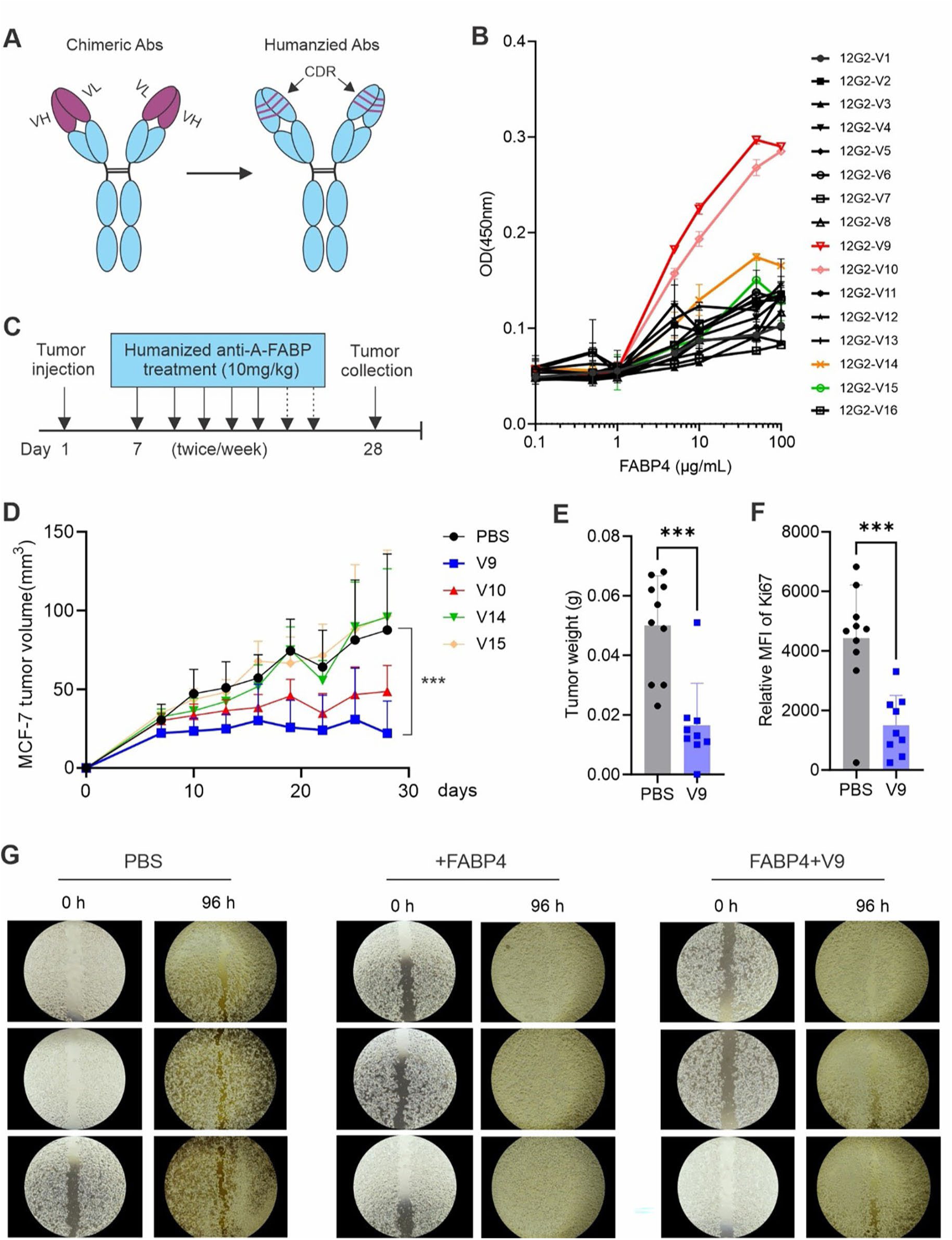
Assessing therapeutic and blocking activity of humanized 12G2 variants. (A) Schematic of production of humanized 12G2 antibody variants. (B) Measurement of 16 humanized 12G2 variants with FABP4 by ELISA (C) Schematic of evaluating the efficacy of selected humanized 12G2 variants using MCF-7 breast cancer model in vivo. (D) Tumor growth curve of MCF-7 in SCID mice treated with V9, V10, V14, V15, or PBS, respectively (***p<0.001). (E) Comparison of tumor weight of MCF-7 in SCID mice treated PBS or V9 for 4 weeks (***p<0.001). (F) Analysis of proliferation of MCF-7 cells by Ki67 expression in SCID mice treated with PBS or V9, respectively (***p<0.001). (G) In vitro analysis of MCF-7 cell migration in the presence of PBS, FABP4 (100ng/ml) or FABP4+V9, for 96 hours.

### Development of humanized 12G2 antibodies

To further humanize the 12G2 antibody, we first aligned the VH and VL sequences of 12G2 mAb with human germline sequences and selected the closest matched germline sequences. The complementary determining regions (CDRs) of 12G2 mAb were then grafted onto the selected human germline sequences with point mutations around the framework amino acids to create 16 humanized 12G2 antibody variants (Figure 3A). We purified the 16 recombinant antibody variants (Figure S3A) and measured their binding affinity to FABP4. Variant 9, 10, 14, 15 showed the highest binding affinity to FABP4 (Figure 3B). Using *in vitro* tumor wound healing assays, we demonstrated that FABP4-enhanced tumor migration could be blocked by these variants, especially by V9 (Figure S3B). Using *in vivo* MCF-7 tumor models, we further demonstrated that compared to other variants, the V9 antibody showed the best therapeutic efficacy in inhibiting MCF7 tumor growth (Figure 3D), tumor weight (Figure 3E) and tumor proliferation (Figure 3F). Consistently, soluble FABP4 enhanced MCF-7 tumor cell migration compared to the PBS control. However, treatment with V9 successfully blocked the FABP4-mediated effect (Figure 3G), supporting a specific role of the V9 antibody.

### Confirming the therapeutic efficacy of V9 antibody using different mouse models

Our previous studies demonstrated that the deficiency of FABP4 inhibited E0771 tumor growth in syngeneic mouse models(18). To compare the efficacy of the V9 antibody, we compared E0771 tumor growth in WT mice treated with or without V9 and in FABP4-/- mice. Tumor growth in mice treated with V9 antibody was significantly slowed down compared to those treated with the PBS control. Interestingly, the tumor growth rate in V9-treated mice was similar to that in FABP4-/- mice (Figure 4A), suggesting that V9 antibody treatment exhibits an equal effect to FABP4 knockout. Moreover, E0771 tumor size and weight in V9-treated mice were similar to those in FABP4^-/-^ mice (Figure 4B, 4C). Consistent with our previous observations in FABP4-/- mice, V9 treatment reduced tumor stemness, as evidenced by reduced ALDH1 activity (Figure 4D). V9 treatment also reduced the production of IL-6 but not TNFα in tumor-associated macrophages compared to PBS-treated mice (Figure S4A-S4D).

**Figure 4.**
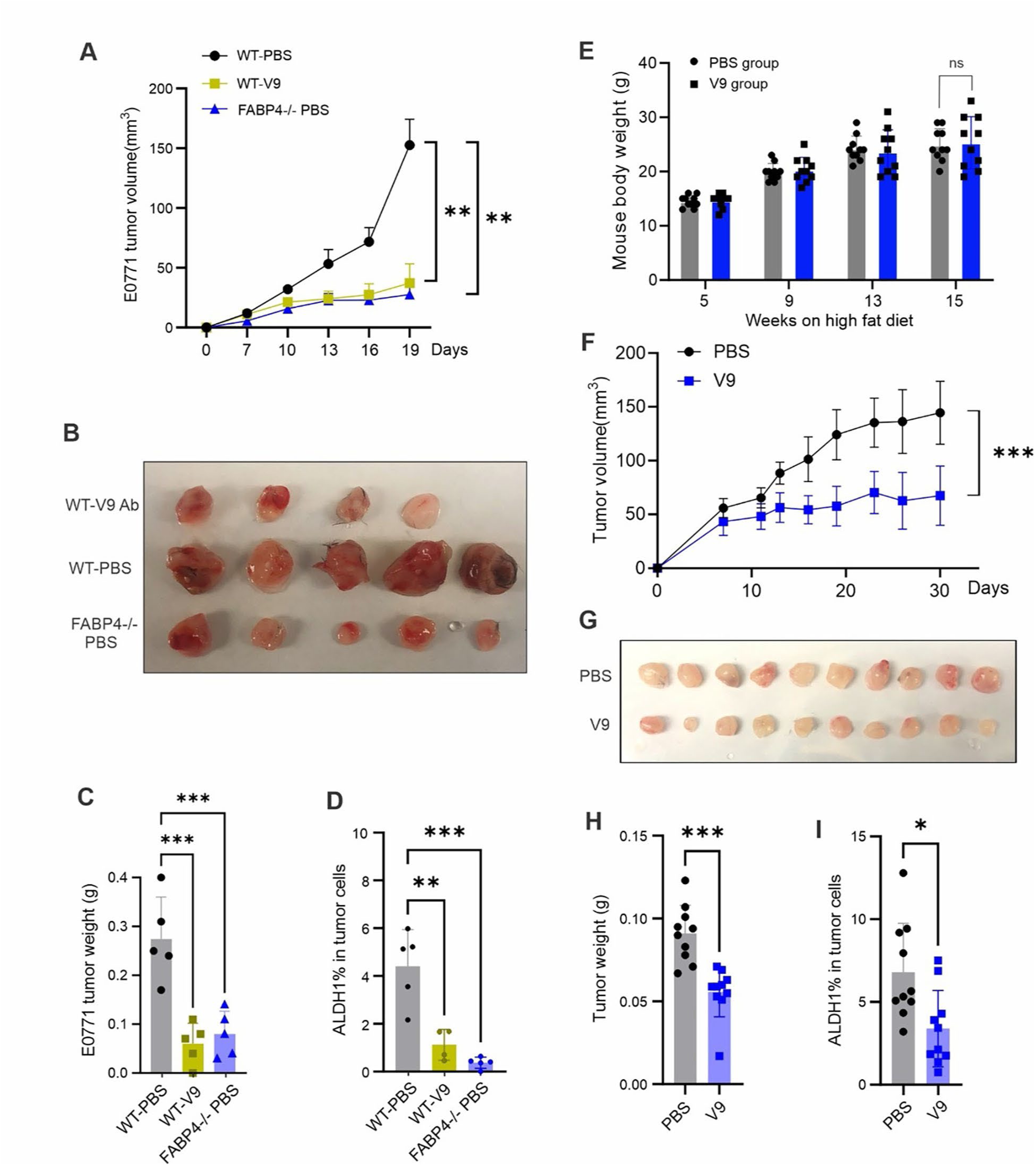
Validation of the efficacy of humanized V9 antibody using different mouse models. (A) E0771 tumor growth curve in WT mice treated with PBS or V9 mAb (10mg/kg) respectively. Tumor growth in FABP4^-/-^ mice treated with PBS was used as a control (**p<0.01). (B) E0771 tumor size in WT mice treated with PBS or V9 mAb and in FABP4-/- mice treated with PBS. (C) E0771 tumor weight in WT treated with PBS or V9 mAb or in FABP4^-/-^ mice treated with PBS (***p<0.001). (D) Analysis of ALDH1 activity in E0771 tumors in mice treated with PBS or V9 mAb (**p<0.01, ***p<0.001). (E) Body weight of SCID mice fed with a HFD for 15 weeks (ns, non-significant). (F) MCF-7 tumor growth curve in HFD-fed SCID mice treated either with V9 mAb (10mg/kg) or PBS control (****p<0.0001). (G-H) MCF-7 tumor size G) and weight (H) in HFD-fed SCID mice treated with V9 mAb or PBS, respectively (***p<0.001). (I) Analysis of ALDH1 activity for MCF-7 tumors in HFD-fed SCID mice treated with PBS or V9 antibody (p<0.05).

Obese mice exhibited elevated levels of circulating FABP4(17). To test the efficacy of V9 antibody in obese mice, we fed SCID mice a high fat diet for 15 weeks, and randomly grouped and treated them with V9 antibody or vehicle PBS after the implantation of MCF-7 tumors in these mice (Figure 4E). Consistently, the efficacy of V9 antibody treatment in obese mice was evident in the reduced rate of tumor growth (Figure 4F), tumor size (Figure 4G), tumor weight (Figure 4H) and reduced ALDH1 activity in tumor cells (Figure I). Notably, when we treated Balb/c mice implanted with the highly aggressive 4T1 mammary tumor cells with the V9 antibody, we did not observe significant tumor growth inhibition (Figure S4E, S4F), suggesting that the efficacy of V9 was not universal for all types of breast cancer.

### Characterization of the V9 antibody

To facilitate the translational potential of V9 antibody, we generated a stable cell pool of V9 antibody (S-V9) using Chinese hamster ovary (CHO) cells and characterized its binding properties of S-V9 antibody. First, we determined the dissociation constant (K_D_) of S-V9 against FABP4 via surface plasmon resonance (Biacore). Kinetic analysis of S-V9/FABP4 interaction showed that S- V9 had an overall K_D_ of 2.07×10^-7^μM with FABP4 (Figure 5A). To delineate the precise binding epitopes of S-V9 on FABP4, we synthesized 25 peptides, each consistent of 15 amino acids (AA) with an overlap of 10 AA to cover the whole 132 AA sequence of FABP4 (Figure 5B). These peptides were biotinylated at the N-terminus to enable binding measurement (Table S). We demonstrated that peptides 1, 9, 11, and 18 exhibited strong binding to S-V9 (Figure 5C), mapping the interaction epitopes to the β-1, β-2/3, β-3/4, and β-7 stand of the FABP4 structure (Figure 5D, Figure S5A), respectively. The patten of epitope recognition by S-V9 distinguished it from other previous FABP4-targeting antibodies, CA33 and HA3 (Figure S5B), underscoring its potential for unique blocking mechanisms in therapeutic applications.

**Figure 5.**
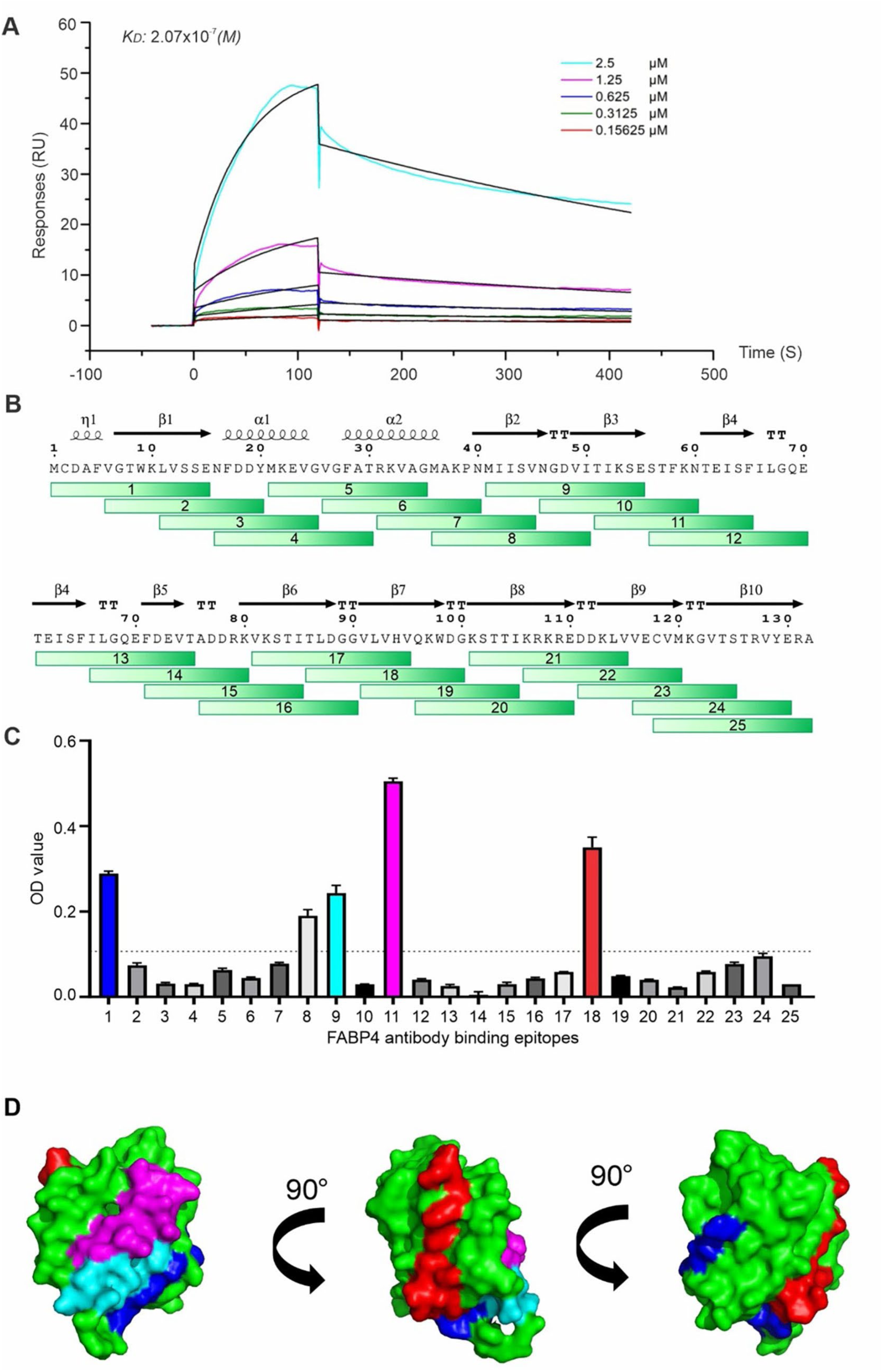
Measurement of V9/FABP4 binding properties. (A) Analysis of K_D_ of V9/FABP4 interaction via SPR technology (Biacore 8K). (B) Synthesis of 25 overlapping peptides spanning the entire length of the FABP4 protein for epitope identification. (C) Measurement of V9 mAb binding peptides in FABP4 by ELISA (D) Different angels of space-filling representation of FABP4/V9 mAb binding generated using the SWISS-MODEL.

### S-V9 mAb inhibits tumor growth and metastasis by disrupting mitochondrial energy metabolism

Using E0771 tumor models, we confirmed the efficacy of the S-V9 mAb in significantly inhibiting E0771 tumor growth, reducing tumor weight and decreasing tumor ALDH1 activity, as consistently observed (Figure 6A-6C). The long-term therapeutic potential of S-V9 was further evaluated by treatment of E0771 tumor-bearing mice over a period exceeding six weeks. Notably, S-V9 treatment resulted in a robust inhibition of tumor growth (Figure S6A-S6B). In PBS-treated group, all mice developed lung metastasis, while half of S-V9 treated mice did not exhibit any lung metastasis (Figure S6C). The metastasis tumor nodules and nodular areas were significantly smaller in S-V9-treated mice compared to those in PBS-treated mice (Figure 6D, 6E). Using Visium spatial single cell transcriptomic analysis (10X Genomics), we found that E0771 tumors exhibited 4 clusters using unsupervised KNN clustering at a resolution of 0.1 (Figure S6D).

**Figure 6.**
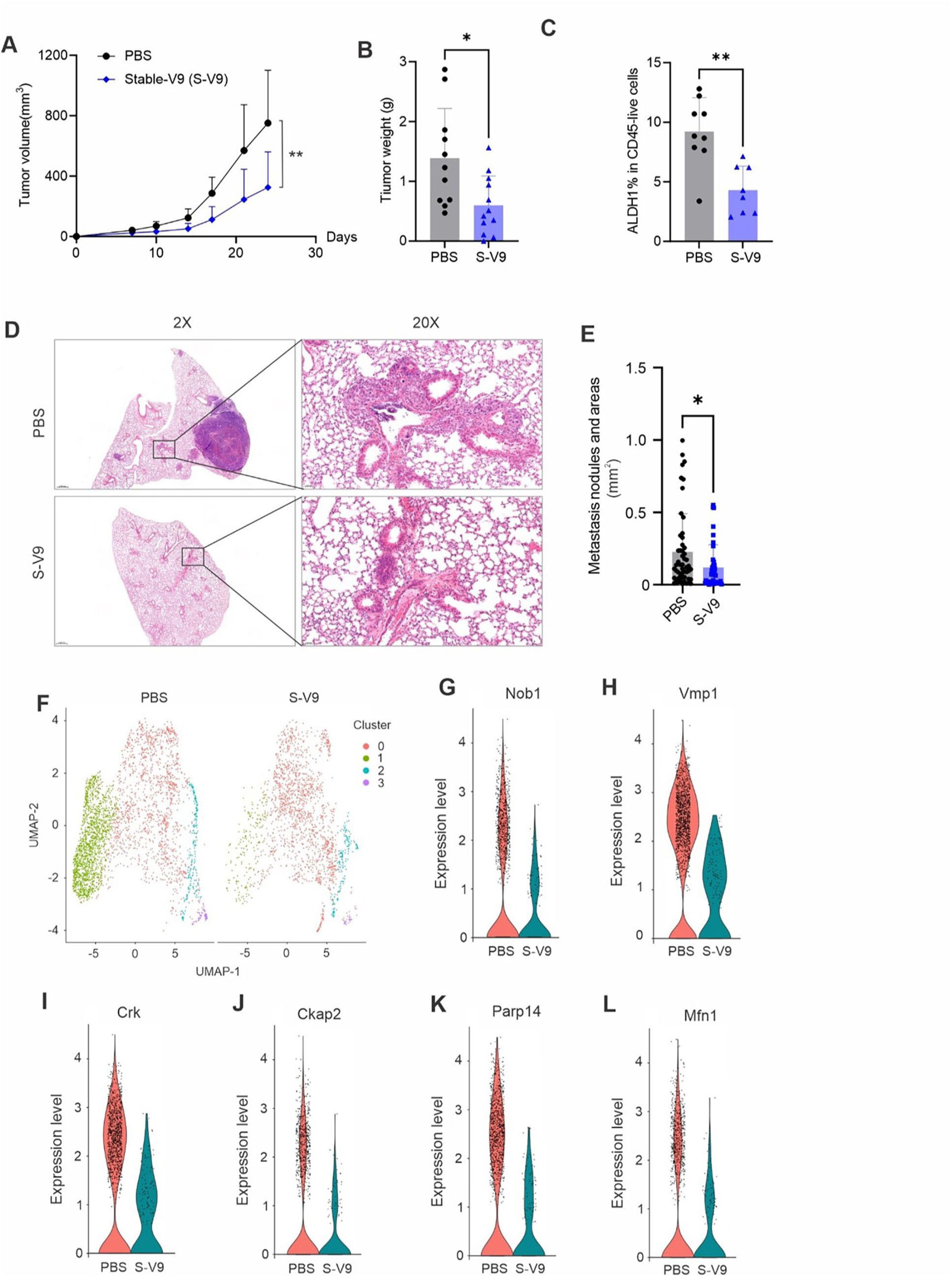
S-V9 antibody treatment inhibits tumor growth and metastasis through inducing abnormal mitochondrial metabolism in tumor cells. (A) E0771 tumor growth curve in mice treated with PBS or S-V9 mAb (5mg/kg) or PBS for 24 days (**p<0.01). (B) E0771 tumor weight on 24 days post tumor implantation in mice treated with S-V9 mAb or PBS (*p<0.05). (C) Analysis of ALDH1 activity in E0771 tumors in mice treated with S-V9 mAb or PBS, respectively (**p<0.01). (D-E) Analysis of lung metastasis of E0771 tumor cells in mice treated with PBS or S-V9 mAb for 24 days by H&E staining. Metastatic nodule numbers and areas are shown in panel E (*p<0.05). (F) UMAP of unsupervised clusters in tumors treated with PBS or S-V9 mAb by spatial transcriptome analysis. (G-L) Violin plot of breast cancer cell markers, including Nob1 (G), Vmp1 (H), Crk (I), Ckap2 (J), Parp14 (K), and Mfn1 (L), on cluster 1 of tumors from mice treated with PBS or S-V9 mAb.

Remarkably, cells in cluster “1” were greatly reduced in response to antibody treatment while the proportion of cells in cluster “0” increased after treatment (Figure 6F, Figure S6E). Among the top 10 marker genes in cluster “0”, we found multiple immune cell markers, including IL1b, CCL4, S100a8, suggesting as an overall immune cell population for the cluster “0”. In cluster 1, breast cancer cell markers, including Nob1, Vmp1, Crk, Ckap2, Parp14 and Mfn1, detected among the top cluster markers (Table S). Expression of these cancer marker genes in cluster “1” were significantly reduced owing to antibody treatment (Figure 6G-6L), suggesting that S-V9 antibody treatment reduced cancer cell aggressiveness. Further analysis of differential expressed genes (DEG) in cluster “1” indicated that antibody treatment mainly affected pathways related to oxidative phosphorylation, mitochondrial protein-containing complexes, electron transport chain and ATP synthesis (Figure S6F). gProfiler human disease phenotypic analysis showed that S-V9 antibody treatment induced abnormal activity of mitochondrial respiratory chain and metabolism (Figure S6G). iPathwayGuide (AdvaitaBio) disease analysis also showed the top two hits to cluster “1” DEGs in response to antibody treatment were “Mitochondrial complex 1 deficiency” and “Combined oxidative phosphorylation deficiency”, further confirming the previous analysis (Table S). Collectively, these results suggested that blocking FABP4 activity with the S-V9 antibody exerts a profound effect on mitochondrial energy metabolism within tumor cells, potentially contributing to the observed therapeutic outcomes.

## DISCUSSION

Based on the expression pattern of hormone receptors (HR) and human epidermal growth factor receptor 2 (HER2) on cancer cells, breast cancer generally falls into four subtypes as luminal A, luminal B, HER2^+^ and triple-negative (25, 26). As such, hormone or HER2-targeted therapies (such as Herceptin, anti-HER2 antibody) are used to treat luminal or HER2^+^ patients. Due to the lack of receptors, triple-negative breast cancer represents the most difficult subtype to treat(27). Given emerging roles of dysregulated lipid metabolism in cancer growth and metastasis(28, 29), blocking FABP4 activity represents a novel strategy for breast cancer treatment as it blocks lipid transportation and metabolism, thus inhibiting cancer cell growth and metastasis.

Bristol Myers-Squibb has developed a small molecular inhibitor, BMS309403, that binds FABP4 and inhibit its function(30). Interestingly, mice treated with BMS309403 exhibited reduced symptoms of diabetes, atherosclerosis and mammary tumor growth(17, 31), suggesting that targeting FABP4 with small-molecular inhibitors might represent a promising approach for treating obesity-associated diseases. However, like many other small-molecule drug candidates, *in vivo* applications of the BMS309403 demonstrated off-target activities(32, 33), and severe side-effects, including suppressing cardiac contractile function(34). To eliminate the potential concerns related to small-molecule inhibitors, highly specific anti-FABP4 antibodies were developed to block the activity of circulating FABP4 in obese models(35, 36). A rabbit polyclonal anti-FABP4 antibody was shown to reduce circulating FABP4 and to improve glucose tolerance in high fat diet-induced obese mice(35). Subsequently, a rabbit-derived monoclonal anti-FABP4 antibody was developed with a therapeutic effect for treating type 2 diabetes in obese mice(36). Notably, lean mice or humans exhibited relatively low levels of circulating FABP4, but tumor-bearing mice or humans exhibited elevated levels of circulating FABP4 suggesting that tumors mobilized adipose tissue lipolysis for their growth benefits(17). These studies suggest that targeting circulating FABP4 with monoclonal antibodies offers a promising strategy for blocking FABP4- mediated effects in both lean and obese subjects.

Although anti-FABP4 antibodies developed in rabbits showed treatment efficacy in animal studies, these non-human antibodies cannot be used in clinical trials or for clinical treatment due to their immunogenicity in humans. To reduce immunogenicity without losing the antibody/antigen-specific binding property, we developed the first humanized anti-FABP4 antibodies in the current studies. As an evolutionarily conservative protein, FABP4 shares high homology between animals and humans(20). To expand the antibody repertoire targeting effective epitopes of FABP4, FABP4 knockout mice were used for human FABP4 immunization in the current studies. After screening over 1200 hybridoma clones, we identified multiple anti-FABP4 monoclonal antibodies. Upon sequencing these antibodies, mouse/human chimeric antibodies consisting of mouse variable domains with human IgG1 constant region domain were developed. Of note, 12G2 clone exhibited a significant therapeutic efficacy by inhibiting tumor growth in a range of syngeneic and xenograft mouse models. To further improve the therapeutic and clinical potential, we generated 16 humanized 12G2 antibody variants by grafting its CDRs into the closest-matching human framework sequences. After assessing their blocking function using *in vitro* cellular studies, we identified that the 12G2-V9 variant exhibited the most effective efficacy by inhibiting mammary tumor growth in various mouse models.

Given that most cancer death is due to metastasis(37), we further demonstrated that V9 treatment significantly inhibited tumor lung metastasis. Using spatial transcriptome analysis, we showed that V9 treatment induced abnormal mitochondrial respiration in tumor cells, contributing to tumor cell death. Epitope analysis of V9 binding to FABP4 indicated that V9 uniquely bound to β1, β-3/4, and β-7, potentially affecting the fatty acid enter and exit of FABP4, thus reducing lipid utilization and metabolism in tumor cells.

During our studies, there were several observations that are worth noting: 1) there were multiple anti-FABP4 mAbs which showed higher binding affinity to FABP4 than 12G2, but high affinity did not always translate into high efficacy in tumor treatment. Antigen-binding epitopes seem to be more relevant to the FABP4 blocking activity *in vivo*. 2) 12G2 inhibited tumor growth in multiple mouse models, including E0771, MMT, MCF-7, but did not appear to be effective for treatment of 4T1 model. it is likely that blocking lipid metabolism by 12G2 does not affect tumor cells (e.g., 4T1 cells) that prefer glucose and glutamine for their rapid cell proliferation(38). 3) Using the doses ranging from 5-30mg/kg in different mouse models, we did not notice any obvious side effects, such as mouse death, reduced body weight, cytokine storm or local skin reactions. We are therefore optimistic that severe adverse effects in future clinical applications will be minimal. Moreover, given the pathogenic role of FABP4 in obesity-associated diseases, including diabetes, atherosclerosis and other cardiovascular diseases.(39, 40), the humanized anti-FABP4 antibodies we generated in this study are expected to have broader applications beyond breast cancer treatment.

## Conclusions

In summary, we identified FABP4 as a new adipokine promoting breast cancer development and developed the first humanized anti-FABP4 monoclonal antibodies for treating breast cancer in mouse models. Blocking circulating FABP4 with monoclonal antibodies represents a novel therapeutic strategy for treatment of breast cancer.

## Abbreviations

FABP4: Fatty acid binding protein 4
aP2: Adipocyte protein 2
IGFs: Insulin/insulin-like growth factors
CDRs: Complementary determining regions
mAb: Monoclonal antibody
S-V9: Stable cell pool of V9 antibody
CHO: Chinese hamster ovary cells
K_D_: Dissociation constant
HR: Hormone receptors
HER2: Human epidermal growth factor receptor 2

## Acknowledgments

The authors would like to thank Iowa Institute of Human Genetics: Genomics Division and Iowa NeuroImaging Processing Core for providing 10x Genomics Visium Spatial Transcriptomics Service.

## Author contributions

Conceptualization: B.L. Methodology: J.H., R.J., Y.Y. X.J., Z.X, and N.S. Spatial RNA sequence analysis: M.C. Writing—original draft: B.L. Writing—review and editing: S.S. Supervision: B.L. Funding acquisition: B.L., S.S. All authors read and approved the final manuscript.

## Data availability

All data needed to evaluate the conclusions in the paper are present in the paper and/or the Supplementary Materials.

## Declarations

### Ethics approval and consent to participate

All experiments were performed according to the approval by the institutional review board at the University of Iowa, Iowa City, IA. Mouse experiments were performed according to protocols approved by the Institutional Animal Care and Use Committee at the University of Iowa, Iowa City, IA.

### Consent for publication

Not applicable

## Supplementary Information

**Table S1.** Selected monoclones specific binding to FABP4.

| Clones | Affinity to FABP4 | Affinity to FABP5 | Ascite production |
| --- | --- | --- | --- |
| 3B10-1 | +++ | - | Yes |
| 3B10-2 | +++ | - | no |
| 3B10-3 | +++ | - | few |
| 5A9-1 | +++ | - | no |
| 5A9-2 | +++ | - | few |
| 5A9-3 | +++ | - | no |
| 6D5 | ++ | - | no |
| 6F12 | + | - | few |
| 9G4-1 | +++ | - | Yes |
| 7A12 | + | - | few |
| 12G2 | ++ | - | Yes |
| 12G7 | ++ | - | few |
| 12A3 | + | - | no |
| 12B1-2 | + | - | few |
| 12B4-1 | ++ | - | few |
| 12H12 | ++ | - | Yes |
| 9D4 | ++ | - | Yes |
| 1A2 | ++ | - | few |
| 1F6 | + | - | few |
| 3C4 | + | - | few |
| 3H10 | +++ | - | few |
| 4G4 | +++ | - | few |
| 6H10 | +++ | - | Yes |
| 13F2 | +++ | - | few |
| 13G7 | ++ | - | few |

Binding affinity: +++ best, ++ better, + good, - negative

**Table S2.**
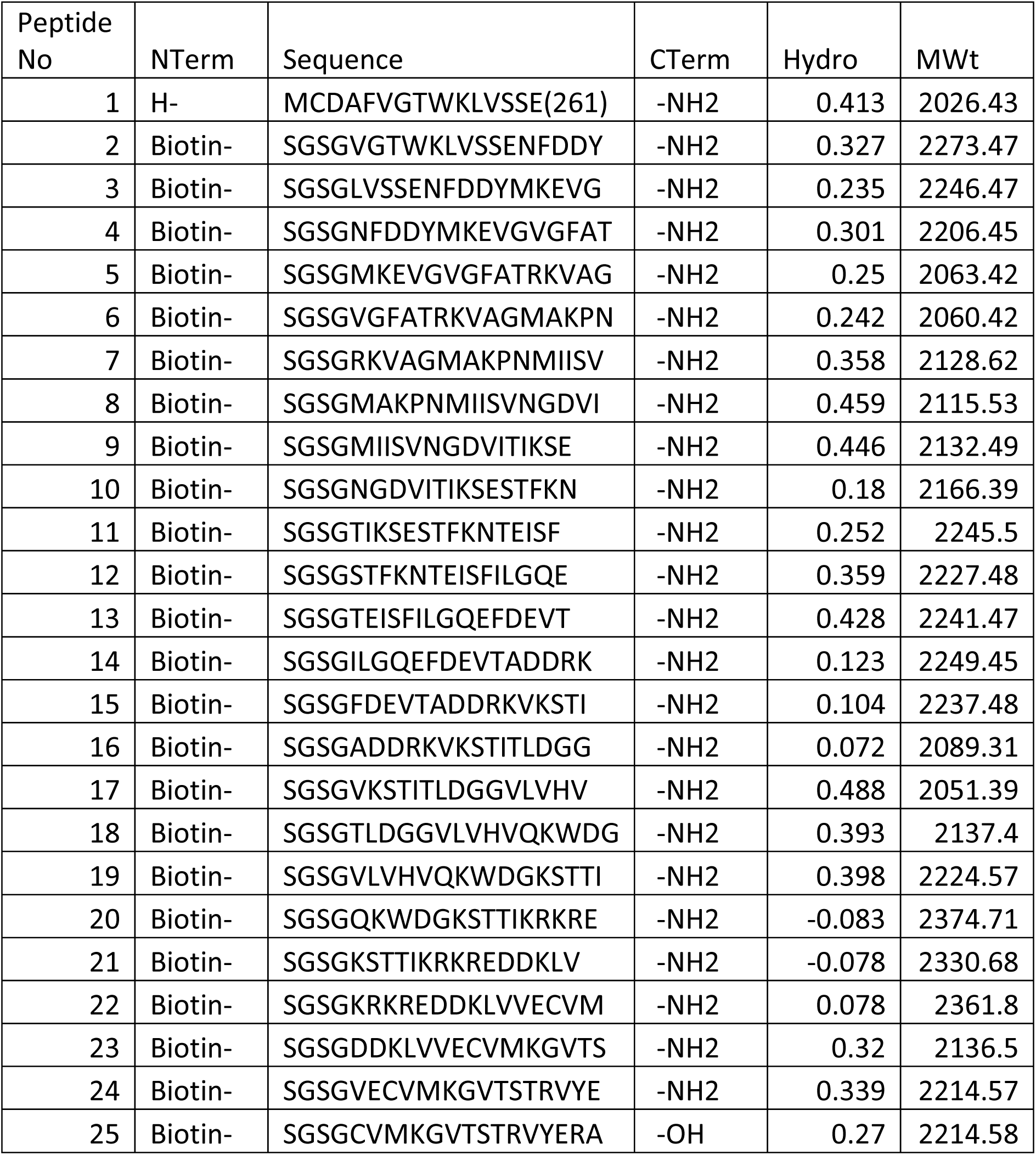
Sythesized biotinylated FABP4 peptides.

**Table S3:** Marker genes of cluster 0 and 1.

| Cluster | p_val | avg_log2FC | pct.1 | pct.2 | p_val_adj | gene |
| --- | --- | --- | --- | --- | --- | --- |
| 0 | 0.00E+00 | 0.9368257 | 0.998 | 0.801 | 0 | Rps13 |
| 0 | 0.00E+00 | 0.9494845 | 0.999 | 0.809 | 0 | Rpl27a |
| 0 | 0.00E+00 | 0.984324 | 1 | 0.91 | 0 | Rps12 |
| 0 | 0.00E+00 | 0.8994034 | 1 | 0.966 | 0 | Fau |
| 0 | 0.00E+00 | 0.9257794 | 1 | 0.983 | 0 | Rpl13 |
| 0 | 0.00E+00 | 0.9487995 | 0.995 | 0.712 | 0 | Mif |
| 0 | 0.00E+00 | 0.8937152 | 1 | 0.826 | 0 | Rpl36 |
| 0 | 0.00E+00 | 1.0102008 | 0.855 | 0.302 | 0 | 1110038B12Rik |
| 0 | 0.00E+00 | 1.099856 | 0.889 | 0.345 | 0 | Gm11808 |
| 0 | 0.00E+00 | 1.192708 | 0.831 | 0.296 | 0 | Fabp5 |
| 0 | 0.00E+00 | 0.94053 | 0.92 | 0.391 | 0 | Tma7 |
| 0 | 0.00E+00 | 0.9457118 | 0.597 | 0.144 | 0 | Lmo1 |
| 0 | 0.00E+00 | 0.9014207 | 0.596 | 0.149 | 0 | Cks2 |
| 0 | 0.00E+00 | 1.1738808 | 0.863 | 0.342 | 0 | Gm10076 |
| 0 | 0.00E+00 | 1.0073564 | 0.557 | 0.129 | 0 | Gm43154 |
| 0 | 0.00E+00 | 0.8968978 | 0.63 | 0.185 | 0 | Snhg8 |
| 0 | 0.00E+00 | 1.3661251 | 0.45 | 0.087 | 0 | Ccl4 |
| 0 | 0.00E+00 | 1.6178051 | 0.415 | 0.075 | 0 | Mmp12 |
| 0 | 0.00E+00 | 0.9512724 | 0.485 | 0.125 | 0 | Ube2a |
| 0 | 0.00E+00 | 1.06225 | 0.369 | 0.06 | 0 | Spink2 |
| 0 | 0.00E+00 | 0.9106443 | 0.426 | 0.093 | 0 | Il1b |
| 0 | 0.00E+00 | 0.9022906 | 0.462 | 0.122 | 0 | Hilpda |
| 0 | 0.00E+00 | 0.9589917 | 0.468 | 0.127 | 0 | Adm |
| 0 | 0.00E+00 | 0.97771 | 0.377 | 0.072 | 0 | Pdcd2 |
| 0 | 0.00E+00 | 0.9866003 | 0.342 | 0.055 | 0 | Ttc32 |
| 0 | 0.00E+00 | 0.944532 | 0.414 | 0.1 | 0 | Lsm12 |
| 0 | 0.00E+00 | 1.0298167 | 0.438 | 0.117 | 0 | Bcl2a1b |
| 0 | 0.00E+00 | 0.9484294 | 0.315 | 0.048 | 0 | Rhox1 |
| 0 | 0.00E+00 | 0.9566189 | 0.393 | 0.096 | 0 | Churc1 |
| 0 | 0.00E+00 | 1.1162666 | 0.334 | 0.101 | 0 | S100a8 |
| 1 | 0.00E+00 | 3.23305 | 0.897 | 0.85 | 0 | Ivns1abp |
| 1 | 0.00E+00 | 2.8148569 | 0.95 | 0.942 | 0 | Ewsr1 |
| 1 | 0.00E+00 | 2.6653159 | 0.996 | 0.999 | 0 | Rpl5 |
| 1 | 0.00E+00 | 2.890204 | 0.998 | 0.999 | 0 | Rpl4 |
| 1 | 0.00E+00 | 2.613298 | 0.767 | 0.84 | 0 | Vmp1 |
| 1 | 0.00E+00 | 2.652628 | 0.698 | 0.774 | 0 | Vps4b |
| 1 | 0.00E+00 | 2.5648621 | 0.672 | 0.72 | 0 | 2410004B18Rik |
| 1 | 0.00E+00 | 2.5937116 | 0.638 | 0.72 | 0 | Crk |
| 1 | 0.00E+00 | 2.6665169 | 0.618 | 0.694 | 0 | Pck2 |
| 1 | 0.00E+00 | 3.0906692 | 0.546 | 0.563 | 0 | Parp14 |
| 1 | 0.00E+00 | 2.8084533 | 0.544 | 0.59 | 0 | Mark3 |
| 1 | 0.00E+00 | 3.4168765 | 0.449 | 0.374 | 0 | Csprs |
| 1 | 0.00E+00 | 2.79001 | 0.531 | 0.594 | 0 | Leo1 |
| 1 | 0.00E+00 | 2.5821561 | 0.492 | 0.606 | 0 | Txndc15 |
| 1 | 0.00E+00 | 2.5530415 | 0.443 | 0.558 | 0 | Tm7sf3 |
| 1 | 0.00E+00 | 2.5693818 | 0.439 | 0.552 | 0 | Eif2b3 |
| 1 | 0.00E+00 | 2.6409359 | 0.425 | 0.528 | 2E-07 | Smc3 |
| 1 | 0.00E+00 | 2.8426031 | 0.382 | 0.437 | 4E-07 | Sh2d5 |
| 1 | 0.00E+00 | 2.5883418 | 0.428 | 0.551 | 2.21E-05 | Ryk |
| 1 | 0.00E+00 | 2.5907065 | 0.419 | 0.54 | 0.00017 | Nob1 |
| 1 | 0.00E+00 | 2.6181509 | 0.411 | 0.518 | 0.000173 | Pqlc3 |
| 1 | 0.00E+00 | 2.6546552 | 0.379 | 0.456 | 0.000479 | Mfn1 |
| 1 | 1.00E-07 | 2.793744 | 0.355 | 0.416 | 0.003516 | Ckap2 |
| 1 | 8.00E-07 | 2.5480239 | 0.387 | 0.486 | 0.024677 | Srek1 |
| 1 | 1.50E-06 | 2.7564053 | 0.35 | 0.416 | 0.049061 | Srpk2 |
| 1 | 1.60E-06 | 2.5842248 | 0.397 | 0.515 | 0.052952 | Csnk1g3 |
| 1 | 4.60E-06 | 2.8382481 | 0.312 | 0.35 | 0.149621 | Gtpbp2 |
| 1 | 1.07E-05 | 2.7692026 | 0.351 | 0.428 | 0.344586 | Acadm |
| 1 | 3.13E-05 | 2.6569848 | 0.366 | 0.462 | 1 | Dnajc21 |
| 1 | 5.53E-05 | 2.6253768 | 0.365 | 0.462 | 1 | Insl6 |

**Table S4:** Diseases associated to differentially expressed genes (DEGs) in cluster 1 by iPathwayGuide analysis.

| Disease Name | countDE | countAll | pv |
| --- | --- | --- | --- |
| Mitochondrial complex I deficiency | 10 | 42 | 7.77E-13 |
| Combined oxidative phosphorylation deficiency | 8 | 61 | 2.38E-08 |
| Hypomyelinating leukodystrophy; Pelizaeus-Merzbacher disease (PMD) | 5 | 34 | 6.10E-06 |
| Dilated cardiomyopathy | 5 | 52 | 5.09E-05 |
| Viral myocarditis | 3 | 10 | 5.22E-05 |
| Dermatitis herpetiformis | 3 | 10 | 5.22E-05 |
| Vogt-Koyanagi-Harada syndrome; Vogt-Koyanagi-Harada disease; Uveomeningoencephalitic syndrome | 3 | 10 | 5.22E-05 |
| Hashimoto thyroiditis | 3 | 11 | 7.13E-05 |
| Graves disease | 3 | 11 | 7.13E-05 |
| Eosinophilic granulomatosis with polyangiitis; Churg-Strauss syndrome | 3 | 11 | 7.13E-05 |
| Multiple sclerosis | 3 | 12 | 9.45E-05 |
| Sjogren syndrome | 3 | 12 | 9.45E-05 |
| Adult onset Still disease; Adult Still disease | 3 | 12 | 9.45E-05 |
| Giant cell arteritis; Temporal arteritis | 3 | 13 | 0.000122 |
| Celiac disease | 3 | 13 | 0.000122 |
| Mohr-Tranebjaerg syndrome | 2 | 3 | 0.000176 |
| Jensen syndrome; Opticoacoustic nerve atrophy | 2 | 3 | 0.000176 |
| Primary central nervous system lymphoma | 3 | 15 | 0.000192 |
| Tuberculosis | 3 | 16 | 0.000235 |
| Asthma | 3 | 18 | 0.000339 |
| Systemic lupus erythematosus | 3 | 20 | 0.000468 |
| Allograft rejection | 3 | 21 | 0.000543 |
| Systemic sclerosis; Systemic scleroderma | 3 | 23 | 0.000714 |
| Oocyte maturation defect | 3 | 24 | 0.000812 |
| Mitochondrial complex III deficiency | 2 | 9 | 0.002052 |
| Type 1 diabetes mellitus | 3 | 35 | 0.002466 |
| Dyskeratosis congenita | 2 | 11 | 0.003102 |
| Congenital symmetric circumferential skin creases; Kunze-Riehm syndrome; Michelin tire baby syndrome | 2 | 11 | 0.003102 |
| Congenital fibrosis of the extraocular muscles | 2 | 13 | 0.004355 |
| Cleft lip and/or cleft palate | 2 | 15 | 0.005804 |
| Keratosis linearis with ichthyosis congenita and sclerosing keratoderma; KLICK syndrome | 1 | 1 | 0.007694 |
| P14 deficiency | 1 | 1 | 0.007694 |
| Ethylmalonic encephalopathy | 1 | 1 | 0.007694 |
| Autoinflammation lipodystrophy and dermatosis syndrome;<br>Proteasome associated autoinflammatory syndromes (PRAAS); Chronic<br>atypical neutrophilic dermatosis with lipodystrophy and elevated<br>temperature (CANDLE) syndrome; Joint contractures, muscle atrophy,<br>microcytic anemia, and panniculitis-induced lipodystrophy (JMP);<br>Japanese autoinflammatory syndrome with lipodystrophy (JASL) | 1 | 1 | 0.007694 |
| Spondylometaphyseal dysplasia, Sedaghatian type | 1 | 1 | 0.007694 |
| Burn-McKeown syndrome | 1 | 1 | 0.007694 |
| Cerebrocostomandibular syndrome | 1 | 1 | 0.007694 |
| Retinal dystrophy, iris coloboma, and comedogenic acne syndrome | 1 | 1 | 0.007694 |
| Leigh syndrome | 2 | 18 | 0.008329 |
| Noonan syndrome and related disorders | 2 | 19 | 0.009262 |
| Myelodysplastic syndrome | 2 | 19 | 0.009262 |
| Juvenile-onset dystonia | 1 | 2 | 0.015328 |
| Nestor-Guillermo progeria syndrome | 1 | 2 | 0.015328 |
| Baraitser-Winter syndrome | 1 | 2 | 0.015328 |
| MEDNIK syndrome; Erythrokeratoderma variabilis type 3 | 1 | 2 | 0.015328 |
| Amyloidosis, Finnish type; Meretoja syndrome; Amyloid cranial<br>neuropathy with lattice corneal dystrophy | 1 | 2 | 0.015328 |
| Phosphoenolpyruvate carboxykinase deficiency | 1 | 2 | 0.015328 |
| Myelodysplastic/myeloproliferative neoplasms | 2 | 25 | 0.015763 |
| Osteogenesis imperfecta | 2 | 30 | 0.022288 |
| Complex cortical dysplasia with other brain malformations | 2 | 30 | 0.022288 |

**Supplementary Figure 1.**
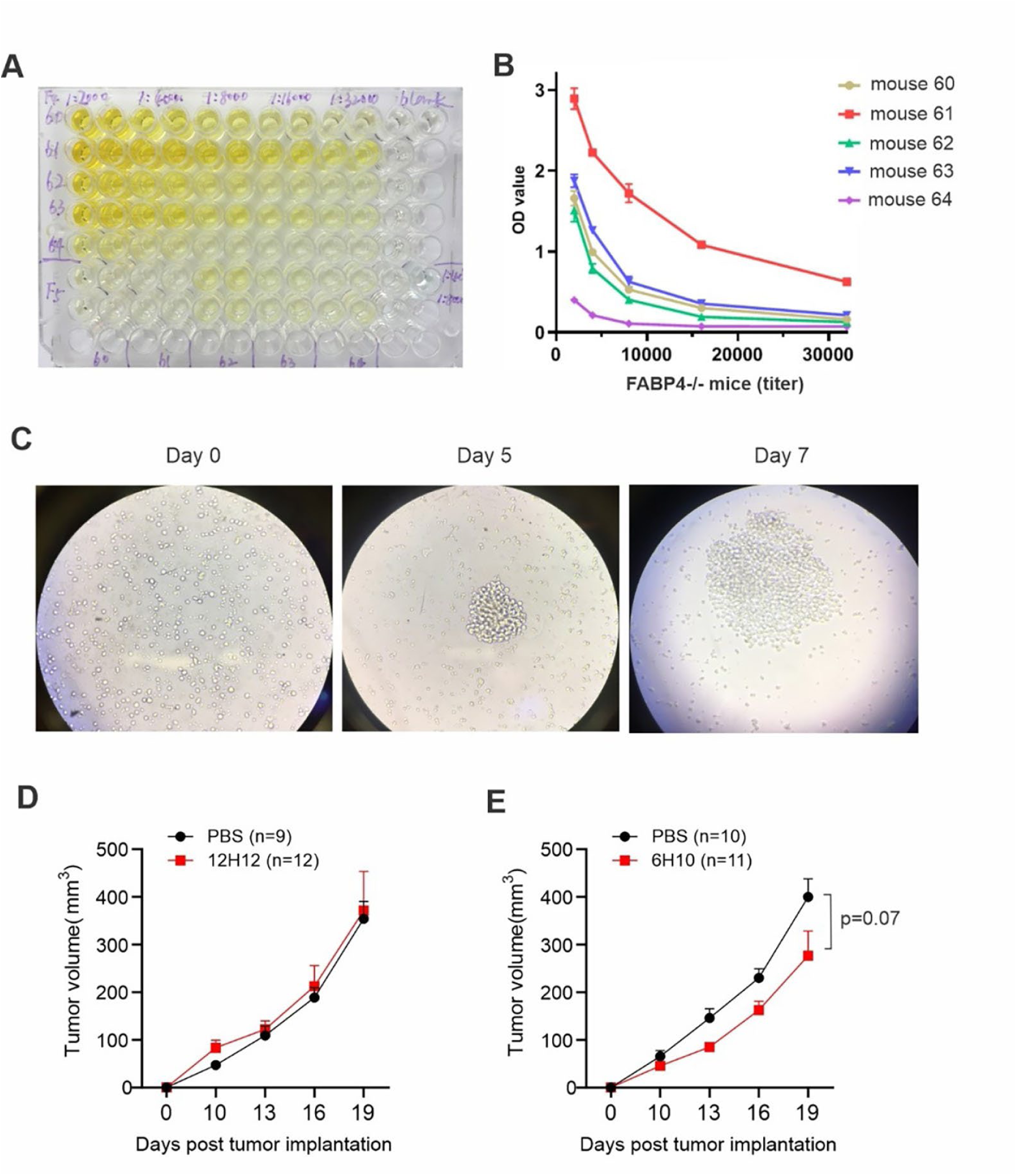
Screening of anti-FABP4 hybridoma clones for the treatment of mammary tumors. (A,B) Evaluating anti-FABP4 serum titer in FABP4-/- mice immunized with recombinant human FABP4 protein. The OD value of anti-FABP4 titer in diluted serum is shown in panel B. (C) Monoclonal hybridoma cells were developed and selected for testing of specific binding to FABP4. (D,E) mAbs from 12H2 (D) and 6H10 (E) clones were purified and used for the treatment of E0771 tumors in mouse models. Antibody treatment (30mg/kg) started from day 7 post tumor injection.

**Supplementary Figure 2.**
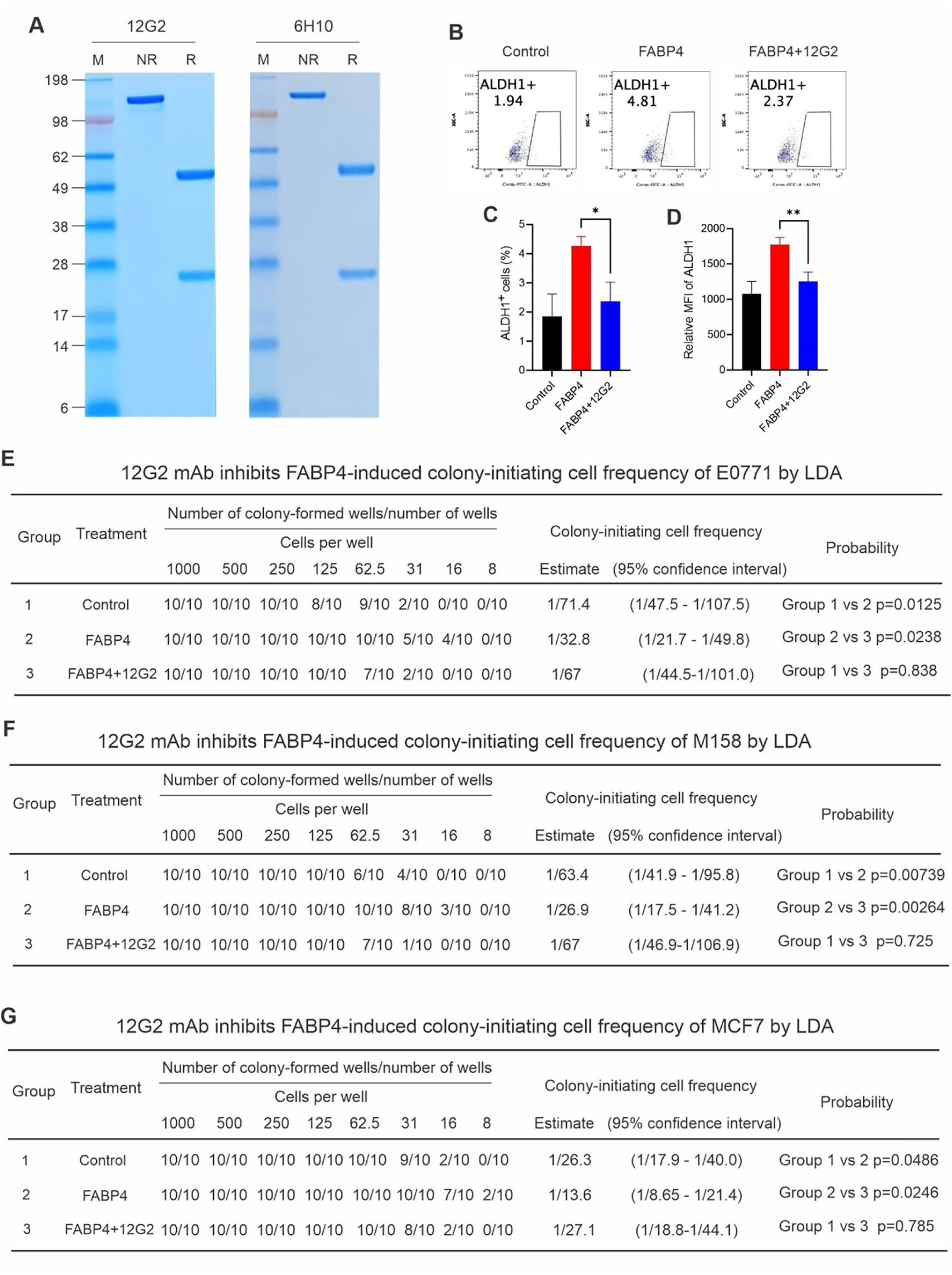
Assessing blocking activity of anti-FABP4 antibodies in vitro. (A) Analysis of purity of the 12G2 and 6H10 antibodies by SDS-PAGE under non-reducing (NR) and reducing (R) conditions. (B-D) MCF-7 cells were treated with PBS control, recombinant human FABP4 (100ng/ml) or recombinant human FABP4+12G2 for 24 hours. ALDH1 activity were measured by flow cytometric staining. Percentage (C) and mean fluorescent intensity (D) of ALDH1+ cells are shown in panel C and D, respectively (*p<0.05, **p<0.01). (E-G) Limiting dilution assays were used to determine the blocking activity of 12G2 in inhibiting FABP4-induced colony-initiating cell frequency using E0771 cells (E), M158 cells (F) and MCF7 cells (G).

**Supplementary Figure 3.**
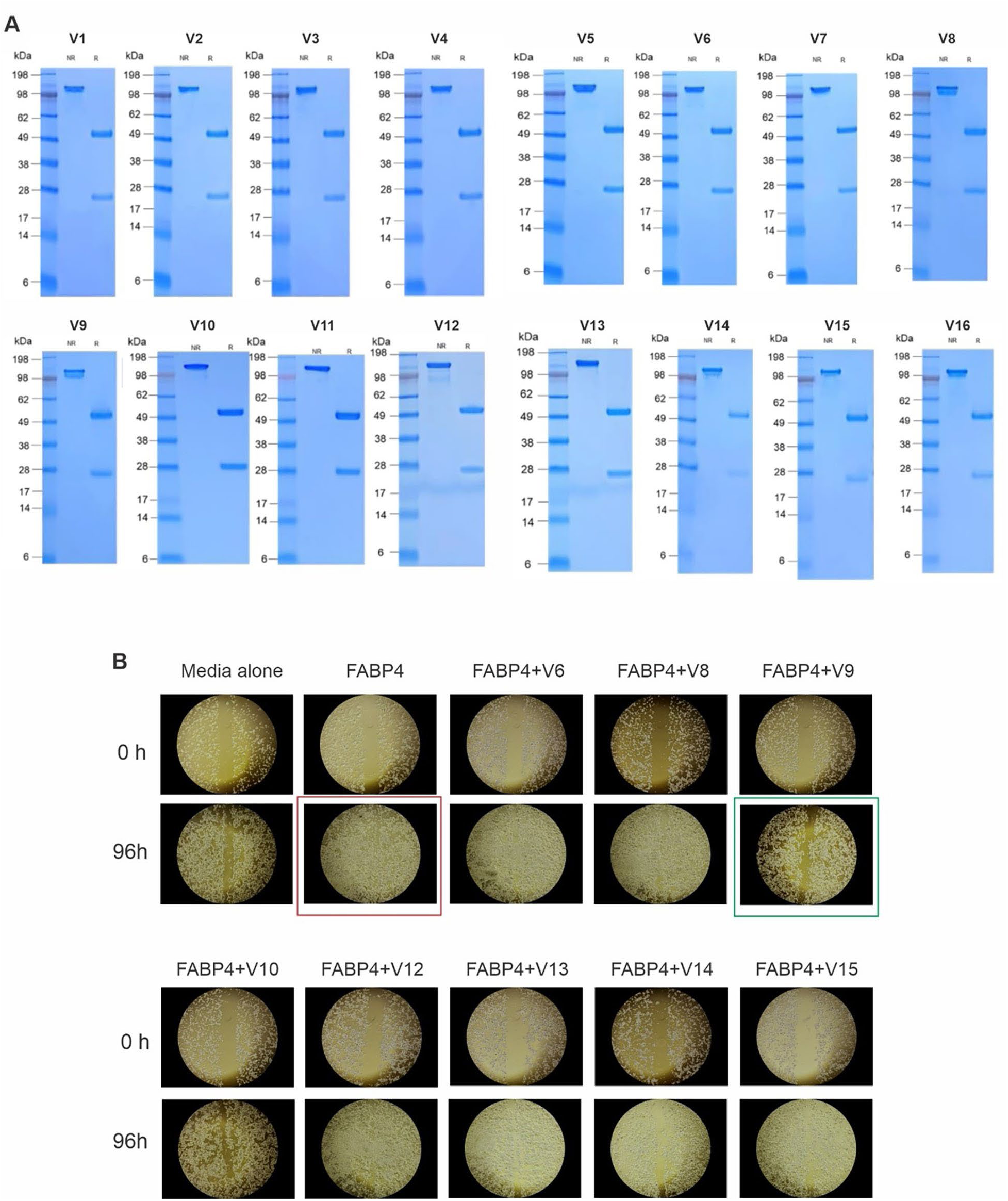
Evaluating the blocking activity of humanized anti-FABP4 antibodies in vitro. (A) Analysis of the purity of 16 humanized 12G2 variants (V1-V16) by SDS-PAGE under non-reducing (NR) and reducing (R) conditions. (B) MCF-7 cells were cultured in medium alone, recombinant human FABP4 alone (100ng/ml) (red square), or recombinant human FABP4+individual humanized antibody variants, for 96 hours. Wound healing assay was conducted to determine that V9 antibody variant (green square) was able to inhibit FABP4-induced MCF-7 migration.

**Supplementary Figure 4.**
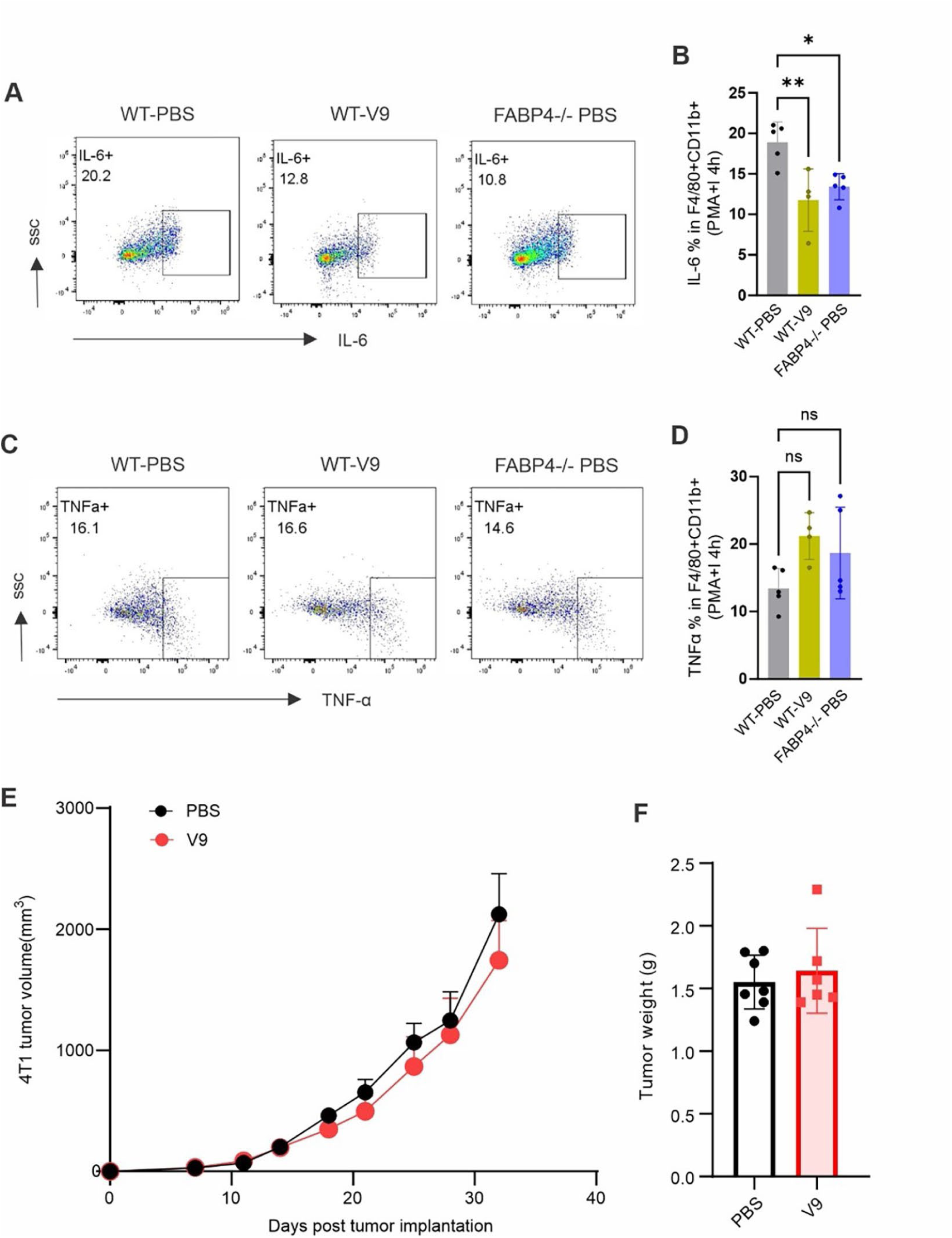
Analysis of the impact of V9 mAb treatment on tumor infiltrating associated macrophages. (A,B) Intracellular staining of IL-6 production in tumor infiltrating macrophages in tumors from mice treated with either PBS or V9 antibody. The percentage of IL-6^+^ macrophages is shown in panel B (*p<0.05, **p<0.01). (C,D) Intracellular staining of TNFα production in tumor infiltrating macrophages in tumors from mice treated with either PBS or V9 antibody. The percentage of TNF-α^+^ macrophages is shown in panel B (ns, non-significant). (E,F) 4T1 tumor growth curve in Balb/c mice treated with PBS or V9 mAb (10mg/kg). Tumor weight on day 33 post tumor implantation is shown in panel F.

**Supplementary Figure 5.**
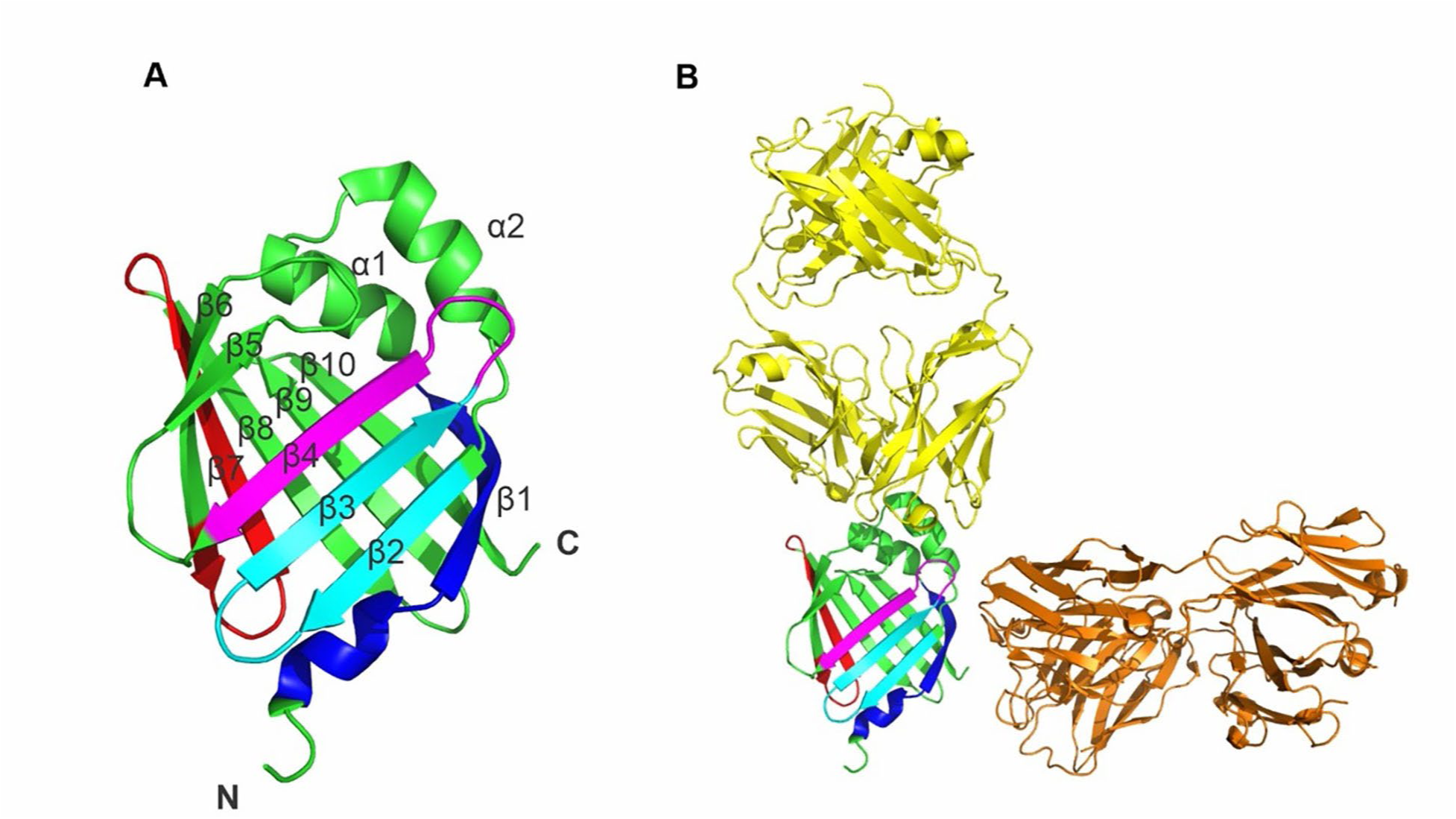
Analysis of V9 mAb binding epitopes within the three dimensional structural of FABP4. (A) Secondary structure image of V9 binding to β-1 (blue), β-2/3 (cyan), β-3/4 (magenta), and β-7 (red) in the structure of FABP4 (PDB 6LJW). (B) Computer modeling of unique binding sites of V9 (magenta) to FABP4 (PDB 5D8J/5C0N) in comparison with other FABP4 antibodies, including CA33 (green) and HA3 (cyan).

**Supplementary Figure 6.**
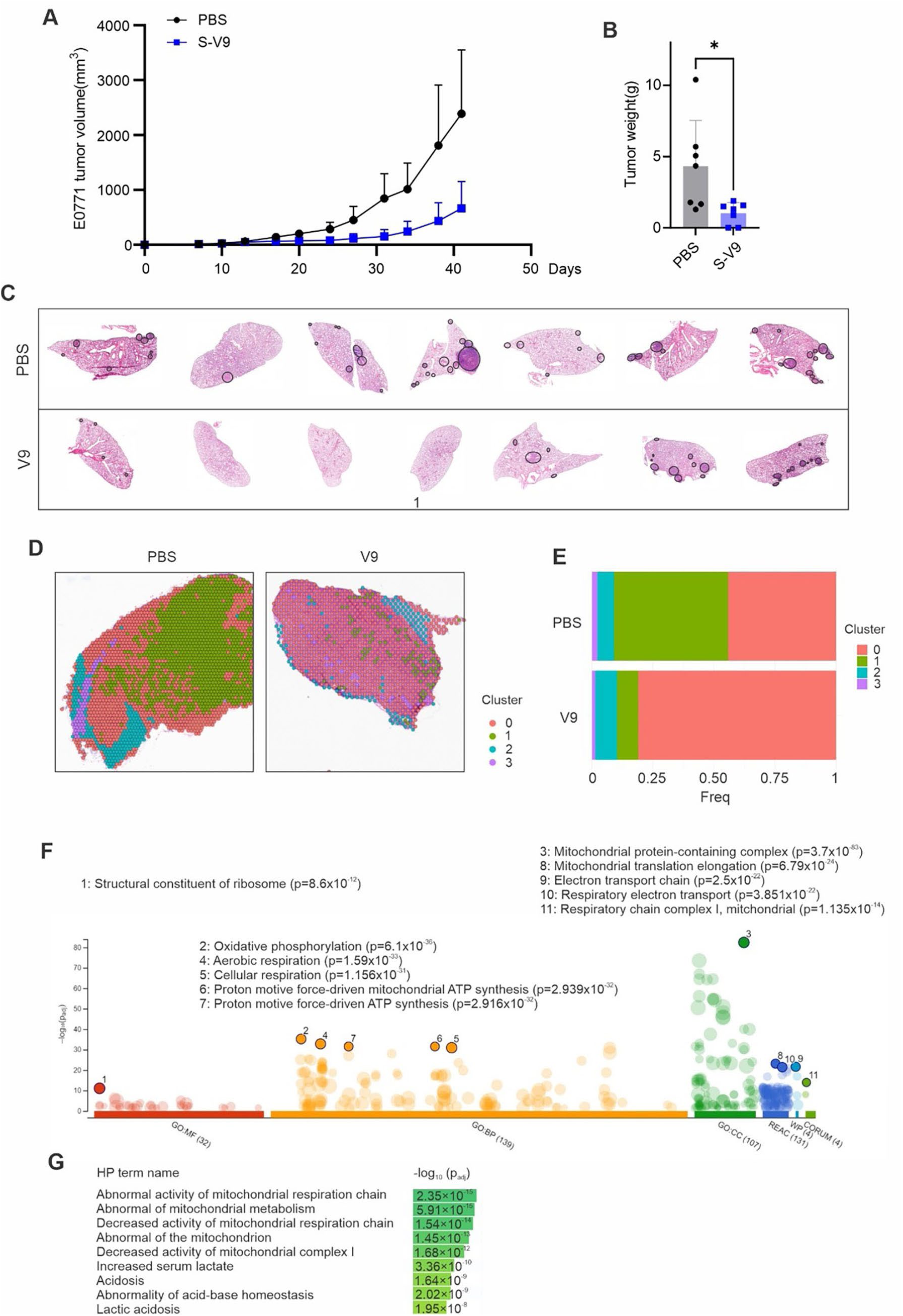
S-V9 mAb treatment inhibited E0771 tumor growth and metastasis. (A-B) E0771 tumor growth curve in mice treated with PBS or S-V9 mAb (5mg/kg) for 6 weeks. Tumor weight is shown in panel B (*p<0.05). (C) Analysis of metastatic nodules (black circle) in lungs of mice treated with PBS or S-V9 mAb by H&E staining. (D) Spatial Dimplot showing unsupervised cell clusters in tumors treated with PBS or S-V9 antibody, respectively. (E) Cluster proportions in tumors treated with PBS or S-V9 antibody, respectively. (F) g:Profiler pathway analysis, including gene ontology (GO), KEGG Reactome (REAC), WikePathways (WP), protein complexes from CORUM using cluster 1 DEGs. (G) g:Profiler analysis of human disease phenotypic (HP) using cluster 1 DEGs

